# Subpopulations of *Pseudomonas aeruginosa* inflict distinct tissue damage during infection

**DOI:** 10.64898/2026.10.06.757000

**Authors:** Alejandro Tejada-Arranz, Varun Sunder, Eduard Basler, Ulrike Lanner, Minia Antelo-Varela, Oliver Biehlmaier, Ridma C. Fernando, Urs Jenal, Alexander Schmidt, Marek Basler

## Abstract

*P. aeruginosa* is a Gram-negative, ubiquitous, opportunistic pathogen that can colonize open wounds and the respiratory airways and is particularly problematic for patients suffering from cystic fibrosis (CF) where it can cause chronic infections. One factor contributing to *P. aeruginosa* diversity during infection is phenotypic heterogeneity. Phenotypic heterogeneity is the ability of bacteria to express different levels of selected pathways, giving rise to multiple bacterial subpopulations in the same environment. The prevalence and relevance of this phenomenon have been poorly characterized in infection models. In this study, we developed strategies to detect these subpopulations via single cell reporters, flow cytometry and fluorescence microscopy, in simple infection models and a physiologically relevant, transwell-based model of the bronchial epithelium. Next, we optimized cell sorting coupled to low input proteomics approaches to characterize the physiological state of the identified subpopulations, and genetically manipulated *P. aeruginosa* to lock them in ON or OFF states of expression of phenotypically heterogeneous pathways. We identify that type IVc/Tad pili are expressed in a subset of *P. aeruginosa* cells and inflict damage to the host tissue, while type Tad pili-negative subpopulations express other virulence factors instead. Lastly, we validated the heterogeneous expression of this machinery across clinical strains. Our findings uncovered the diversity of strategies employed by *P. aeruginosa* for pathogenesis, where distinct bacterial subpopulations use different virulence factors to damage respiratory epithelia.

## Introduction

*P. aeruginosa* is a Gram-negative, ubiquitous, opportunistic pathogen that can colonize open wounds and the respiratory airways, and is particularly problematic for patients suffering from cystic fibrosis (CF)^1^. In these patients, *P. aeruginosa* is able to establish chronic infections that can last decades. During this time, *P. aeruginosa* undergoes a pathoadaptive process where certain mutations are under strong selective pressure^2^, contributing to strain diversification within the airways of infected patients^3,4^. Besides this intrahost evolution, *P. aeruginosa* is known for its metabolic diversity and ample repertoire of virulence factors, being able to grow in a variety of environments and inflict damage to many eukaryotic hosts. One further factor contributing to *P. aeruginosa* diversity during infection is phenotypic heterogeneity. Phenotypic heterogeneity is the ability of bacteria to express different levels of selected pathways, giving rise to multiple bacterial subpopulations in the same environment. This phenomenon has been shown to be relevant for virulence and antibiotic resistance in other pathogens such as *Salmonella enterica*^5–7^. Some studies have described the existence of phenotypic heterogeneity in *P. aeruginosa*, but most studies so far have focused on biofilm or surface colonization settings^8–11^, with limited evidence of the relevance of this phenomenon for the infection process^12,13^.

In this study, we identify multiple pathways that display heterogeneous expression levels in *P. aeruginosa* cells across different infection models. Using cell sorting and low input proteomics, we describe the physiological state of some of these distinct bacterial subpopulations in an air-liquid interface model of the bronchial epithelium. Furthermore, we generated *P. aeruginosa* strains locked in ON or OFF states of expression of selected pathways and identified that distinct *P. aeruginosa* subpopulations inflict different levels of damage to the host tissue and elicit differential responses. Lastly, we identify a previously unrecognized role of Tad/Type IVc pili on tissue damage through cilia destruction.

## Results

### Specific pathways are heterogeneously expressed in infection models

To identify pathways that are differentially expressed in *P. aeruginosa* subpopulations during infection, we constructed a collection of fluorescent, plasmid-based transcriptional reporters based on pathways that were found to be consistently up- or downregulated in infected human samples versus *in vitro* culture conditions^14–16^. We selected 73 promoters driving the expression of pathways important for anaerobic respiration, nutrient acquisition, biofilm lifestyle, anabolic processes, transcriptional regulators and selected virulence factors. The full list of promoters, predicted operon genes and rational for their selection can be found in Supplementary Table 1. The shortlisted promoters were cloned upstream from an *mNeongreen* gene in a pSEVA231 backbone (Supplementary Figure 1A), and the resulting plasmids were introduced into *P. aeruginosa* PAO1. To detect putative promoters driving heterogeneous gene expression due to the metabolic environment of the CF airways, we first measured mNeongreen expression in Synthetic Cystic Fibrosis sputum Medium (SCFM). We screened the reporter collection for bulk expression of mNeongreen in a plate reader, and at the single cell level by flow cytometry during exponential growth (after 4h of incubation) or in stationary phase (after 24h) (Supplementary Figure 1B-C, 2 and 3). Of note, all reporter strains grow similarly well (Supplementary Figure 1B), but display a wide range of fluorescence expression (Supplementary Figure 1C). To identify promoters that drive heterogeneous gene expression upon contact with host cells, we infected A549 alveolar epithelial cells for 3h with the same collection of *P. aeruginosa* strains. Upon lysis of the A549 cells, the fluorescence of single bacteria was analysed by flow cytometry (Supplementary Figure 4). We next calculated whether the resulting distributions of mNeongreen expression followed a unimodal or non-unimodal distribution (Figure 1A) and their bimodality coefficients (Figure 1B). Interestingly, around 25% of the examined reporters displayed heterogeneous behaviours (concomitant non-unimodal distribution with dip p-value<0.05 and bimodality coefficient >0.55) regardless of the examined condition (26% in SCFM after 4h, 23% in SCFM after 24h and 25% on A549 cells after 3h), suggesting that phenotypic heterogeneity spontaneously arises in infection models. Of note, the specific reporters displaying heterogeneous behaviours only partially overlap across conditions, and none are considered by these statistical tests as heterogeneous across all conditions, suggesting that the heterogeneous behaviours are dependent on the specific experimental conditions (Figure 1C).

**Figure 1.**
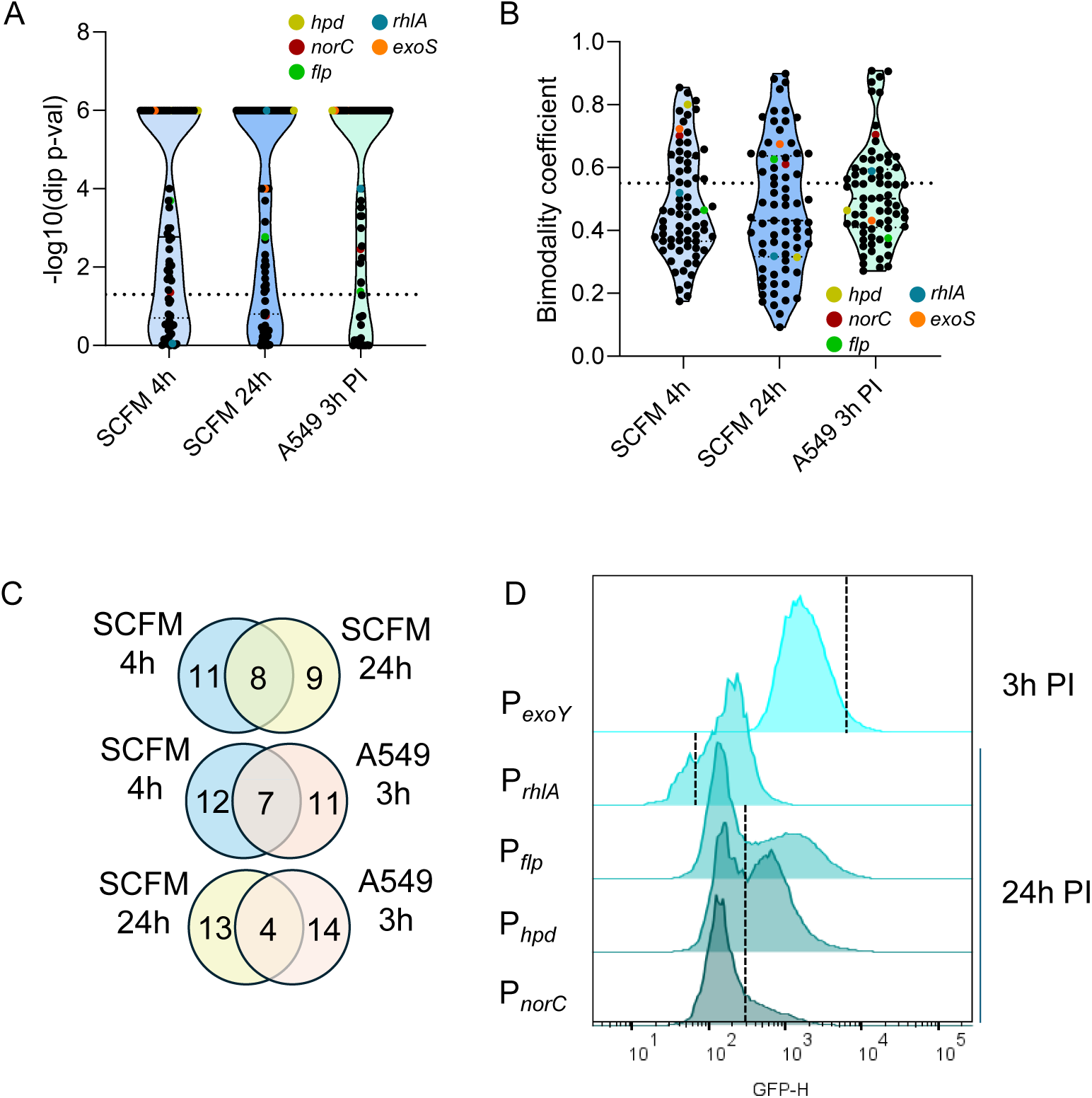
Selected *P. aeruginosa* promoters display heterogeneous expression profiles across conditions. (A) Violin plot of the Hartigan’s dip test statistic across the tested conditions. Each point represents the statistic for a single reporter strain. Highlighted are the reporters selected for further testing in the NHBEC-ALI model. (B) Violin plot of the bimodality coefficient across the tested conditions. Each point represents the statistic for a single reporter strain. Highlighted are the reporters selected for further testing in the NHBEC-ALI model. (C) Venn diagrams showing the overlap of heterogeneously expressed reporters across conditions. (D) Flow cytometry histogram of the Green Fluorescent Protein (GFP) expression profile of selected reporters in the NHBECALI model at the indicated timepoint.

We selected candidate pathways for further analysis that display heterogeneous expression during growth in SCFM (*exoY*, reporter for the expression of the type III secretion system (T3SS); or in stationary phase (*flp*, reporter for the expression of Tad pili).

### Distinct *P. aeruginosa* subpopulations arise during infection of bronchial epithelia

To determine whether these phenotypically heterogeneous subpopulations also arise in more complex infection models, we utilized a Normal Human Bronchial Epithelial Cell (NHBEC) Air-Liquid Interface (ALI) model (from here on, NHBEC-ALI model) that has been used to study infection dynamics by *P. aeruginosa*^17^ and that recapitulates a fully differentiated airway epithelium. We characterized the infection progression of the *P. aeruginosa* strain used in this study on this model (Supplementary Figure 4A). In contrast to the *P. aeruginosa* PAO1 strain used in the previous study (DSMZ lineage), the *P. aeruginosa* PAO1 strain (UW-lineage) used here displays slower tissue disruption dynamics (approx. 2-fold slower tissue breaching), consistently with well-documented genetic and phenotypic differences present across PAO1 strains from different laboratories^18^ (Supplementary Figure 5A) and highlighting the need for phenotype cross-verification across strains. The *P. aeruginosa* PAO1 strain from the UW-lineage (hereafter *P. aeruginosa* PAO1), was subsequently used to infect the NHBEC-ALI model with 1000 CFUs as described to allow for *P. aeruginosa* replication on the tissue. With that strain, tissue breaching by *P. aeruginosa* occurs at around 24h post-infection (hpi), followed by replication in the basal side (Supplementary Figure 5A).

To identify all bacteria in the samples, regardless of the expression status of the reporter of interest, we integrated a constitutively expressed *mCherry2* gene, expressed from the *rplM* promoter in a convergent orientation to *mGreenLantern* (Supplementary Figure 5B). The *rplM* promoter displays strong and unimodal expression across all tested conditions (Supplementary Figure 5C). Importantly, this plasmid backbone is stable in *P. aeruginosa* in the NHBEC model for up to 24h, with some loss at later timepoints (Supplementary Figure 5D) and it does not affect replication on the tissue (Supplementary Figure 5E). Furthermore, the constitutive reporter does not affect the appearance of bacterial subpopulations for the reporters of interest (Supplementary Figure 5F).

We reasoned that the best timepoint to assess the appearance of phenotypic heterogeneity is at the time where tissue destruction starts to become apparent (at 24 hpi), when there are both intact and damaged tissue areas. At 24 hpi, we disrupted the tissues and measured the mGreenLantern fluorescence intensity of bacteria (mCherry2-positive events). The *flp* reporter displayed heterogeneous expression profiles at 24 hpi (Figure 1D). In contrast, the *exoY* reporter, which was previously shown to be induced in a bistable manner, did not do so on the NHBEC-ALI model. This is in contrast to *in-vitro* conditions, where this reporter recapitulates bistable behavior upon addition of the calcium chelator ethylene glycol-bis(β-aminoethyl ether)-N,N,N’,N’-tetraacetic acid (EGTA) (Supplementary Figure 2G). Instead, a non-unimodal expression profile of the *exoY* was identified in the model, indicating that there are still bacteria expressing higher T3SS levels than others (Figure 1D). We conclude that heterogeneity in the gene expression of selected pathways also occurs in more complex models such as the NHBEC-ALI model.

### T3SS-positive subpopulations inflict more damage to the bronchial epithelium and display cell division defects

To isolate different subpopulations of *P. aeruginosa* and characterize their proteome, we optimized a low input proteomics pipeline that allows us to obtain high coverage proteomes from sorted *P. aeruginosa* cells by taking advantage of the recent development of timsTOF proteomics technology^19^. For further information on the optimization process, refer to Supplementary File 1 and Supplementary Figures 6-9. By sorting the 25% of *P. aeruginosa* cells with top and bottom P*_exoY_*-mGL expression at 3 hpi, we could identify an enrichment of T3SS-associated components in the high expression subpopulation (2-fold increase in expression of both T3SS effectors and structural components, Figure 2A and B, and Supplementary Table 2 and 3), indicating that there is a continuum of T3SS expression among *P. aeruginosa* cells that can be identified with our method, regardless of the lack of obvious bistability of the reporter under these conditions. The 3 hpi timepoint was chosen based on previous reports that the T3SS is induced upon contact with host cells^20^. Interestingly, we could observe that some bacterial cells that were T3SS-positive (high mGL) tend to be more elongated at 24 hpi (Figure 2C and D).

**Figure 2.**
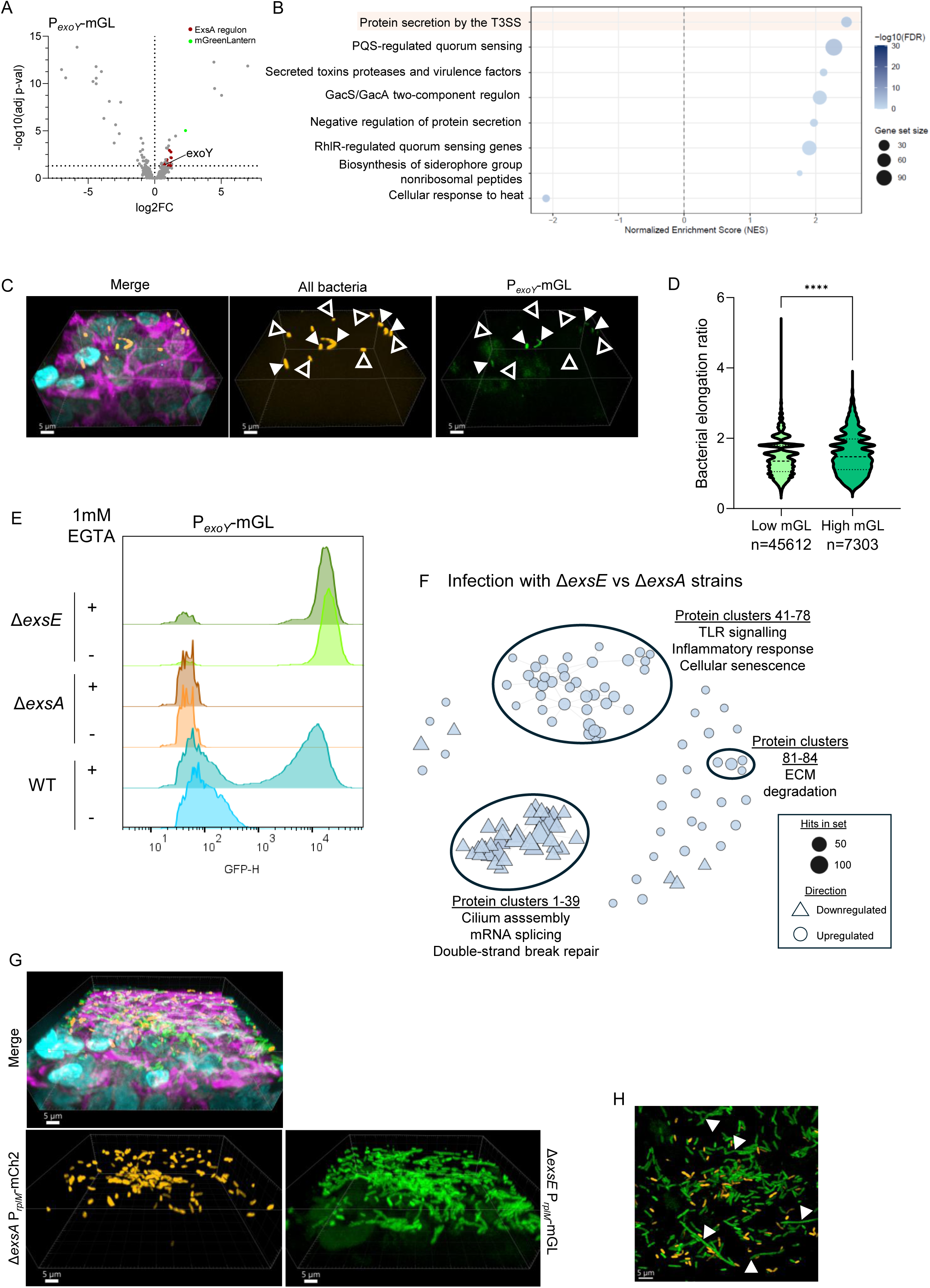
Functional consequences of heterogeneous T3SS expression. (A) Volcano plot of the differences in the proteome of P*_exoY_*-mGL high vs low sorted subpopulations. (B) Gene Set Enrichment Analysis (GSEA) dot plot showing pathways enriched among the dysregulated pathways between high and low of P*_exoY_*-mGL sorted subpopulations. (C) Fluorescence microscopy 3D reconstructions showing heterogeneous T3SS across single bacterial cells. (D) Microscopy-based quantification of the bacterial elongation ratio between high and low P*_exoY_*-mGL bacteria. (E) Flow cytometry histograms of P*_exoY_*-mGL expression across different strains, under standard conditions or after supplementation with 1 mM EGTA (T3SS inducer). (F) Pathway network representation of differentially detected human proteins at 24 hpi with T3SS-ON (Δ*exsE*) or -OFF (Δ*exsA*) *P. aeruginosa* strains. The labels indicated selected identified protein clusters. The full description of the identified clusters and dysregulated hits can be found in Supplementary Table 4. (G) Fluorescence microscopy 3D reconstructions showing co-infecting T3SS-OFF or -ON cells on the tissue surface at 24 hpi. (H) Slice at the tissue surface. Highlighted with white arrows are filamenting T3SS-ON bacteria.

We next constructed deletion mutants with altered T3SS levels. As expected, deletion of the main T3SS transcriptional activator, *exsA*, locked the *P. aeruginosa* cells in an OFF state of expression, while deletion of the repressor *exsE* led to cells locked in an ON state, whether 1 mM EGTA was present or not (Figure 2E). To test how host cells respond to infection with those mutants, we first measured the proteome of WT-infected vs uninfected tissues at 24 hpi. As expected, the human bulk proteome of WT-infected tissues shows signature responses to infection (Supplementary Figure 10, Supplementary Table 4 and 5), such as upregulation of inflammatory response and antigen presentation genes, LPS, IL-1 and IL-17 response genes, as well as genes involved in oxidative phosphorylation, thermogenesis and response to reactive oxygen species (ROS). In addition, we could identify a depletion of microtubule associated genes consistent with a partial destruction of the upper ciliated layer of the tissue. Infection with ON vs OFF-locked bacteria for 24h followed by proteomics indicated that the bacterial subpopulation with increased T3SS activity induces a higher degree of TLR signaling, cellular senescence and inflammatory response, and the induction of the expression of enzymes that degrade the extracellular matrix (ECM) (Figure 2F and Supplementary Table 5 and 6). At the same time, the infected tissues display a depletion of cilium assembly proteins as well as mRNA splicing and double-strand break repair pathways, consistent with a higher degree of tissue destruction. Therefore, we observe the expected impact of T3SS-ON vs OFF, suggesting that even apparently minor skews (around 1 log difference in fluorescence intensity between brightest and dimmest cells using fluorescent reporters) in the expression profile of virulence determinants may have profound consequences for tissue integrity and disease progression.

Interestingly, in a co-infection experiment with T3SS-ON, constitutively mGL-positive and T3SS-OFF, constitutively mCh2-positive strains we could observe frequent bacterial filamentation specifically in cells that were blocked in a T3SS-positive state at 24 hpi (Figure 2G and H). This suggests that cells blocked in a T3SS-positive state display cell division defects, in agreement with our earlier observation that cells with higher T3SS expression tend to be longer than their isogenic, low-T3SS sister cells.

### Tad pili contribute to tissue destruction

Flp is the major pilin of the Type IVc (T4c)/Tad pili, whose function in *P. aeruginosa* has not been explored. The expression of Tad pili is induced upon carbon starvation and stationary phase features by the transcription factor PprB^21,22^. To confirm that *flp* reporter activity correlated with Tad pili protein levels, Flp-positive and -negative bacteria were sorted from tissues at 24 hpi and their proteome was quantified. Indeed, Flp-positive cells display higher level of expression of the components of the Tad pilus, as well as quorum sensing and stationary phase-regulated pathways (Figure 3A and B, Supplementary Table 2 and 7). In agreement with this observation, WT bacteria displayed higher levels of Flp in tissue areas with high bacterial density and lower distance to the nearest bacterium (Figures 3C, D and E). In addition, canonical virulence pathways such as the T3SS and the T4a pili have higher expression in the bacterial subpopulation with low *flp* promoter activity (Figure 3B).

**Figure 3.**
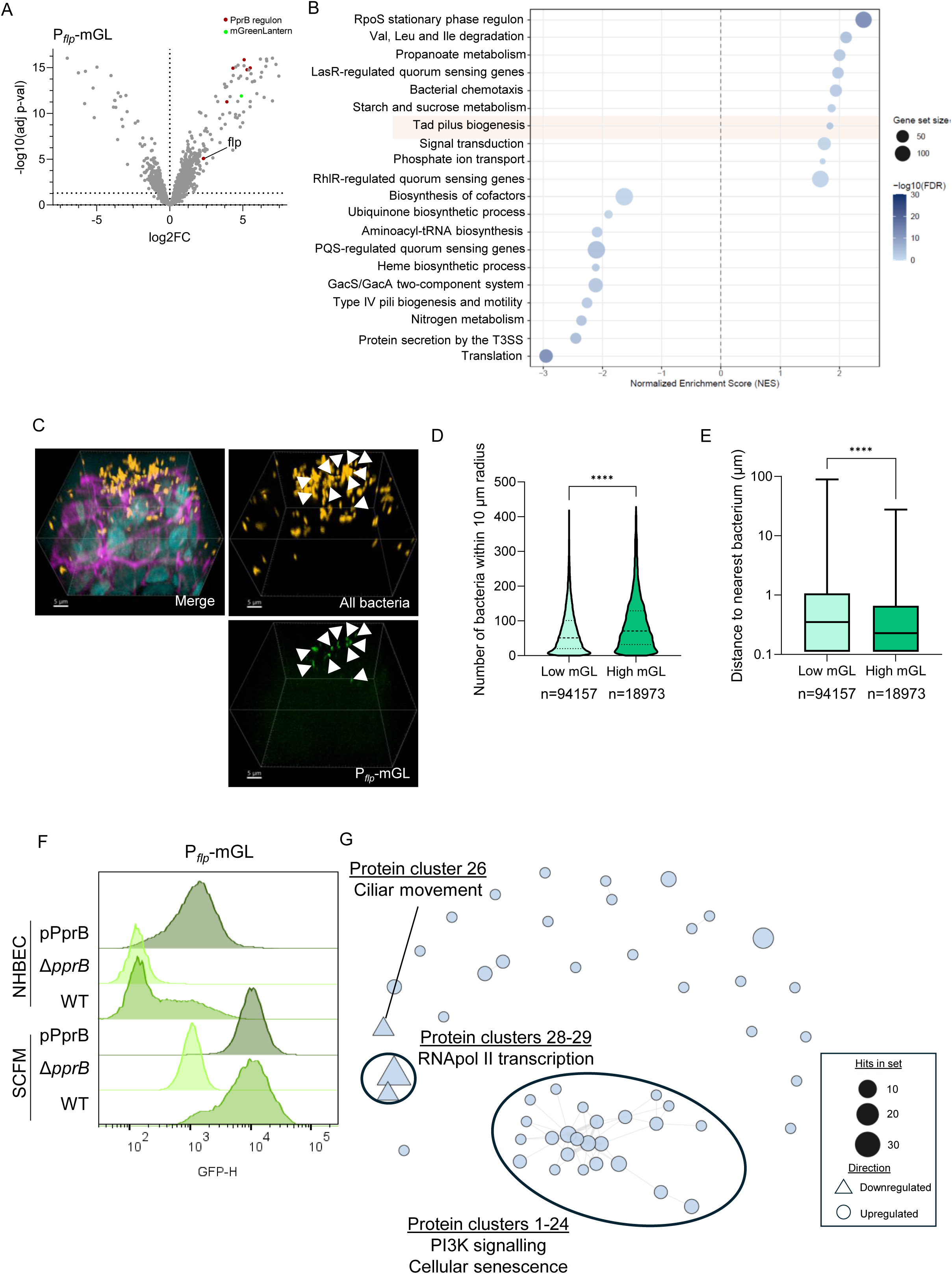
Functional consequences of heterogeneous Tad pili expression. (A) Volcano plot of the differences in the proteome of P*_flp_*-mGL high vs low sorted subpopulations. (B) Gene Set Enrichment Analysis (GSEA) dot plot showing pathways enriched among the dysregulated pathways between high and low of P*_flp_*-mGL sorted subpopulations. (C) Fluorescence microscopy 3D reconstructions showing heterogeneous Flp expression across single bacterial cells. (D) Microscopy-based quantification of bacterial density (number of bacteria within a 10 µm radius) between high and low P*_flp_*-mGL bacteria. (E) Microscopy-based quantification of the distance to the nearest bacterium between high and low P*_flp_*-mGL bacteria. (F) Flow cytometry histograms of P*_flp_*-mGL expression across different strains,in SCFM or in the NHBEC-ALI model at 24 hpi. (G) Pathway network representation of differentially detected human proteins at 24 hpi with Tad pili-ON (pPprB) or -OFF (Δ*pprB*) *P. aeruginosa* strains. The labels indicated selected identified protein clusters. The full description of the identified clusters and dysregulated hits can be found in Supplementary Table 5.

To explore the role of the Tad pili during infection, we constructed strains lacking Tad pili through deletion of the transcription factor *pprB*, or overexpressing Tad pili through supplementation of the WT strain with an extra *in trans* copy of *pprB* (pPprB strain), resulting in bacterial populations locked on OFF or ON states of expression, respectively, in SCFM as well as in the NHBEC-ALI model (Figure 3F). Of note, plasmid pPSV35 used to overexpress PprB has no impact on bacterial growth on the apical side and is also maintained during infection in the absence of selection for at least 30h (Supplementary Figures 11A and B).

We next infected the NHBEC-ALI model with the strains locked on ON or OFF states of expression of Tad pili for 24h and performed bulk proteomics. Despite no significant differences in CFU count at 24 hpi (Supplementary Figure 11C), indicative of comparable growth capabilities of the Δ*pprB* mutant on the airway surface, a lower abundance of bacterial proteins was measured for the Δ*pprB* mutant in our proteomics experiments (Supplementary Figure 11D), likely due to bacterial detachment during the sample preparation process and suggesting a potential attachment defect of this strain. In agreement, 83% less Δ*pprB* bacteria could be detected in the tissue compared to WT by fluorescence microscopy and less apparent tissue destruction (Supplementary Figure 11E and F). On the host side, we could observe a more significant depletion of RNA polymerase II-related transcription factors (1.5-90X depletion) and cilia-associated proteins (1.5-5X depletion), as well as higher levels of cellular senescence markers and increased PI3K signaling when infecting with the pPprB strain (Figure 3G and Supplementary Table 5 and 8), suggesting a further role of Tad pili in attachment to the tissue and tissue destruction.

### Tad pili are heterogeneously expressed across clinical isolates

To confirm whether Tad pili display heterogeneous expression profiles across multiple *P. aeruginosa* strains, we selected eight strains from the previously characterized collection of 414 clinical isolates^23^. According to previously published *in vitro* RNA-seq data, four of these isolates should display high levels of expression of Tad pili genes, and four should display low levels of expression (Figure 4A and B). After introduction of our P*flp* reporter plasmid, we could determine that isolates with expected low levels of expression of Tad pili lack mGL expression in SCFM; whereas isolates with expected high levels of expression of Tad pili display a heterogeneous expression profile similar to that of the PAO1 strain (Figure 4C). We further selected four isolates, two with high and two with low level of *in vitro* expression of Flp and verified their expression in the NHBEC-ALI model at 24 hpi, where the strains show the same expression patterns (high and low, respectively) by flow cytometry and fluorescence microscopy (Figure 4D and Supplementary Figure 12). In addition, in agreement with our previous observations, Tad pili are negatively co-regulated with T4a pili across the collection of clinical isolates (Figure 4E). We propose that Tad pili are expressed in a bistable manner during infection across several *P. aeruginosa* strains, which in turn express lower levels of the canonical T4a pili.

**Figure 4.**
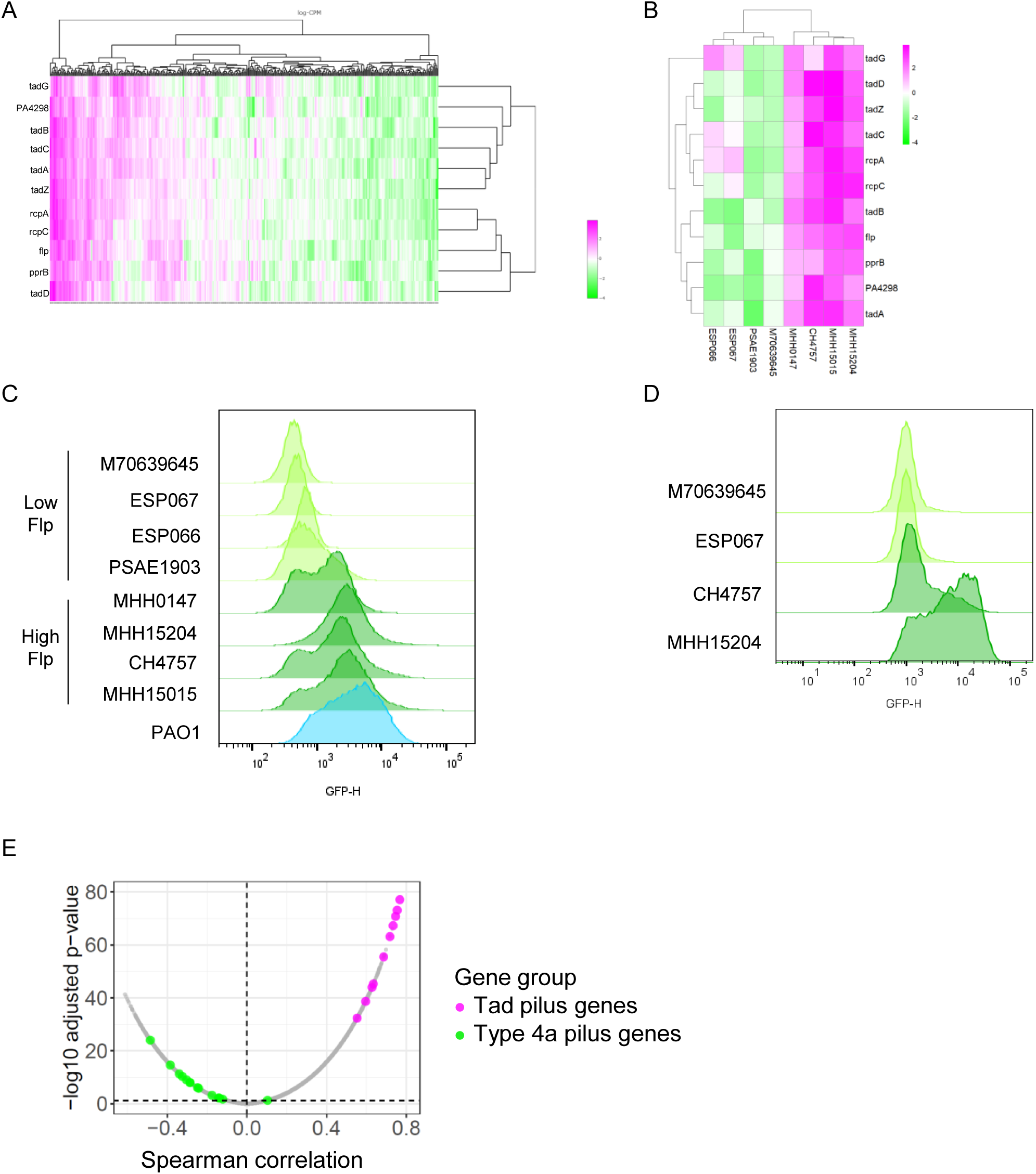
Tad pili are differentially expressed across clinical isolates. (A) Heatmap of the *in vitro* expression level of Tad pili components in LB across clinical isolates. Generated from published RNA-seq data^23^. (B) Heatmap of the *in vitro* expression level of Tad pili components in LB in a subset of 8 selected clinical isolates. (C) Flow cytometry histograms of P*_flp_*-mGL expression in 8 clinical isolates in SCFM. (D) Flow cytometry histograms of P*_flp_*-mGL expression in 4 clinical isolates in the NHBEC-ALI model at 24 hpi. (E) Spearman correlation coefficient of genes that are positively or negatively coregulated with Flp expression. Highlighted are genes encoding components of Tad pili and T4a pili. Generated from published RNA-seq data^23^.

### Tad pili-positive cells attach to cilia and contribute to cilia degradation

From our proteomics dataset, one of the main depleted categories when infecting the tissue with the pPprB versus the Δ*pprB* strain is comprised by cilia-related proteins, suggesting a role of Tad pili in cilia destruction (Figure 5A). To validate this, we stained infected samples with antibodies against ARL13B, a protein that localizes to the ciliary membrane that is structurally essential for motile cilia^24^. By quantifying the intensity of the ciliary signal around WT and Δ*flp* bacteria in a co-infected sample, we could detect that WT bacteria have 2-fold higher ARL13B signal in the neighboring volume (3 µm radius around the bacteria), showing that WT bacteria are more proficient in cilia attachment than Δ*flp* cells (Figure 7B and C) and supporting a role for Tad pili in cilia binding and destruction.

**Figure 5.**
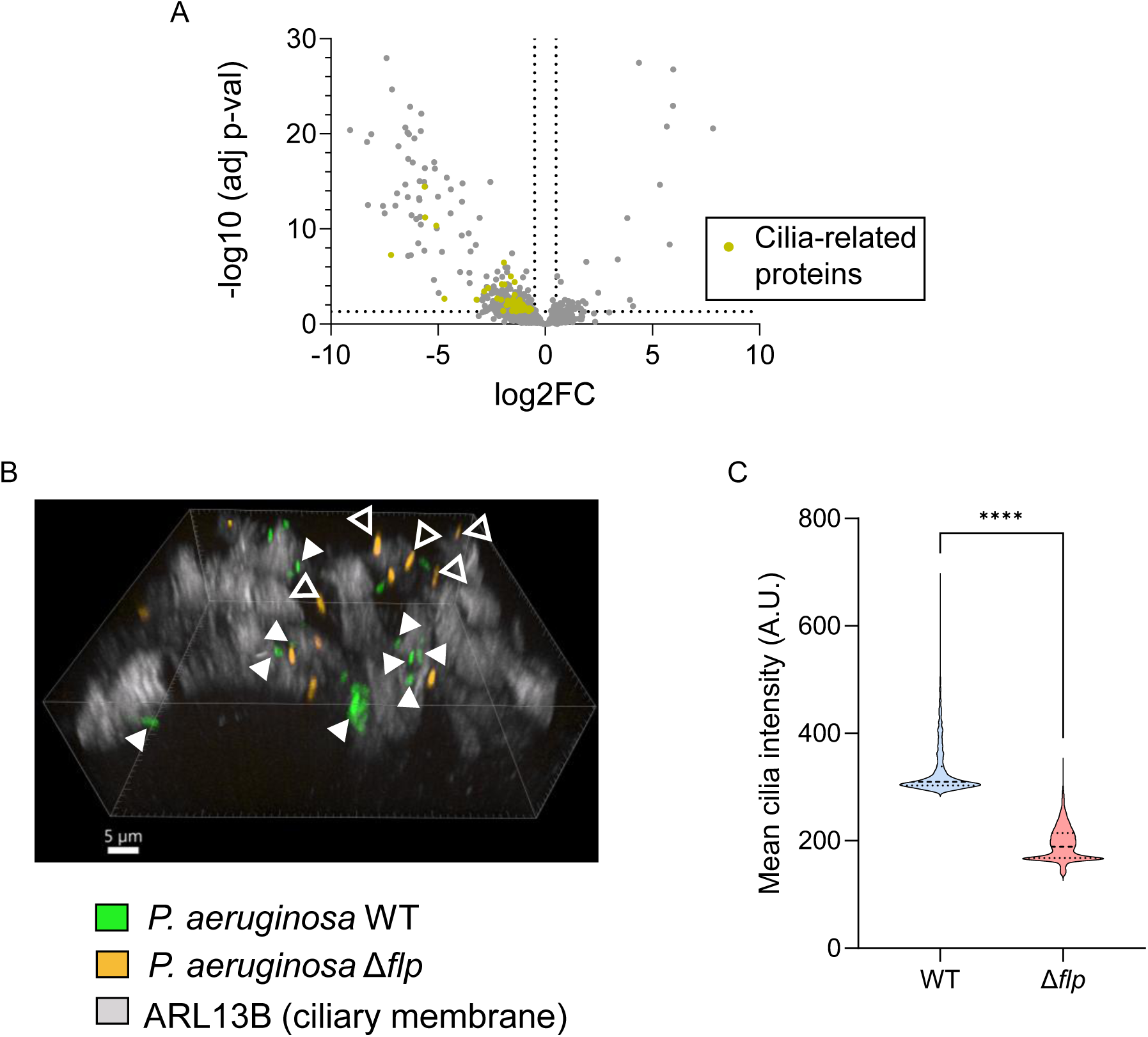
Tad pili support adhesion to cilia. (A) Volcano plot showing differentially detected human proteins at 24 hpi with Tad pili-ON (pPprB) vs Tad pili-OFF (Δ*pprB*) *P. aeruginosa*. (B) Fluorescence microscopy 3D reconstruction of co-infecting WT (green) and Δ*flp P. aeruginosa* (orange) on the tissue surface at 24 hpi. In white, cilia labelled with α-ARL13B antibodies. Highlighted with full white arrows are WT bacteria embedded between the ciliary structures; and in white empty arrows Δ*flp* bacteria in regions with no ciliary signal. (C) Quantification of median cilia intensity within a 3 µm radius from segmented bacteria at 24 hpi.

## Discussion

Although the importance of phenotypic heterogeneity for bacterial populations has been recognized for years^5,25–27^, it has been challenging to describe the properties of the identified bacterial subpopulations and the role they play during infection, due to lack of sensitivity of downstream analysis technologies^28^. Here, we describe a set of tools and approaches to characterize the physiology of bacterial subpopulations and tackle the challenge of assigning a function to these subpopulations.

First, we developed a collection of fluorescent reporter tools to probe the expression profile of selected infection-relevant pathways in *P. aeruginosa* populations. By doing so, we could identify that about 25% of tested reporters display heterogeneous expression profiles under our infection-mimicking conditions. Of note, our selection of the subset of promoters that we tested was biased towards promoters driving the expression of genes that are differentially regulated in human infection samples compared to *in vitro* culture^14–16^. Therefore, we do not claim that 25% of all *P. aeruginosa* reporters display heterogeneous expression profiles across conditions. In another study where 3,025 promoters of *P. aeruginosa* were screened, 42 promoters (1.4%) displayed heterogeneous expression profiles in exponential phase and 115 in stationary phase (3.8%)^29^. However, in this study bacteria were grown in rich medium (LB), and the reporter constructs were screened in a dual colour reporter system. Although we did not systematically test this, we found that a dual colour reporter system may lead to the underestimation of heterogeneously expressed promoters, likely due to transcriptional interference between selected promoter pairs that are dependent on the orientation of the constructs. Therefore, each dual reporter construct needs to be individually validated against single reporter constructs. This, together with the higher complexity of the experimental conditions that we used in this study (infection models) may account for the differences in the apparent prevalence of heterogeneously expressed promoters in *P. aeruginosa.* In contrast, bimodality was found to be rare *in vitro* in *E. coli* ^30,31^, which may reflect key differences in the regulatory strategies employed by these organisms or the relative simplicity of the culture conditions used in those studies.

Besides the few genome-wide studies indicated above, *P. aeruginosa* phenotypic heterogeneity has only been mechanistically characterized for specific pathways, under relatively simple conditions (surface attachment^9–11^, biofilm formation^8,32–35^), with one recent study assessing the importance of one heterogeneously expressed pathway (glycerol metabolism) on virulence against murine macrophages^12^ and the well-known T3SS bistability under inducing conditions (low Ca^2+^, contact with host cells)^20,36^. Our screen identified several pathways that display heterogeneous expression profiles under infection-relevant conditions. Some of them were identified in the studies indicated above (*norC*, *flp*, *exoY*), but others are new (*hpd*, *algU*, *fhp*, *aprX*, among others). Furthermore, we validated the appearance of some distinct subpopulations in the more physiologically relevant NHBEC-ALI model.

Interestingly, we found that, in the NHBEC-ALI model, the expression of the T3SS does not follow the classical bistable-like pattern that has been well described *in vitro*. Instead, WT cells display a narrower range of expression that may suggest that further regulatory mechanisms act in complex infection models to limit T3SS expression. In agreement with this, bacterial cells lacking the repressor *exsE*, that show constitutively high T3SS expression, often display cell division defects, which is in agreement with previous observations reporting longer division times for T3SS-ON cells *in vitro*^37^. Further research is needed to determine why overexpression of the T3SS leads to cell filamentation during infection.

We further characterized the role of the bacterial subpopulations expressing Tad pili during infection. By using flow cytometry, we could identify that around 30% of bacterial cells display high levels of expression of Tad pili during infection of the NHBEC-ALI model. Cells with high P*_flp_*-mGL expression tend to be in areas with higher cell density and closer to their neighbouring bacteria (Figure 5D and E), in agreement with previous observations that Tad pili are upregulated during carbon starvation and stationary phase conditions^21,22^. Furthermore, by manipulating the levels of the transcription factor driving the expression of Tad pili, PprB, we could identify a potential role of Tad pili-positive subpopulations in attachment to the host tissue and cilia destruction.

Although Tad pili have not been directly shown to be regulated by quorum sensing (QS), at least the signals that regulate their expression (carbon starvation, stationary phase) might often co-occur with QS-related signals. In agreement with this, sorted bacterial subpopulations with high Tad pili expression (high P*_flp_*-mGL) display also higher expression levels of QS-regulated factors, as well as amino acid and propanoate degradation genes. Indeed, *P. aeruginosa* cells were also found to display single cell heterogeneity in their levels of expression of QS pathways^38,39^. Interestingly, bacterial populations with low levels of Tad pili (low P*flp*-mGL) display higher levels of other virulence-related factors including the T3SS and the canonical Type IVa pili, highlighting the diversity of virulence strategies used by *P. aeruginosa* populations to inflict damage on the host, where both Tad pili-ON and -OFF populations express important virulence genes.

Another point to consider is strain diversity. Our results show that even PAO1 strains from different sublineages can have vastly different behaviours in complex infection models. This is not surprising, given that large genomic and phenotypic differences have already been reported across PAO1 strains from different laboratories^18^. In this study, we could determine that the PAO1 strain from the UW lineage displays slower tissue destruction dynamics (breaching of the epithelial barrier around 24 hpi) compared to that of the DSMZ lineage (breaching at 9-12 hpi)^17^. However, we could validate that the *flp* promoter displays heterogeneous expression profiles in axenic media and in the NHBEC model across different clinical isolates, in agreement with previously published RNA-seq data^23^. This suggests that there are two classes of *P. aeruginosa* strains according to their *flp* expression level: those where Tad pili are silenced (isolates PSAE1903, ESP066, ESP067 and M70639645 in our study) and those where Tad pili are heterogeneously expressed (isolates MHH15015, CH4757, MHH15204 and MHH0147, as well as PAO1-UW). We could not identify any obvious patterns in the mutations in the promoter or coding sequences of the Tad pili genes or their regulators (*pprB*, *pprA*) between high and low Tad pili strains. In addition, our analysis showing the negative correlation of the expression of Tad and T4a pili *in vitro* suggests that there may be a condition-dependent expression switch between the two types of pili, with different isolates favouring the expression of one type of pilus over the other. Further research is needed to examine the functional consequences of this preference, and the molecular mechanisms triggering Tad pili expression in a subset of cells. Some possibilities may be differential immunogenicity of these pili machineries, or escape from phage predation. While a wealth of phages target T4a pili for bacterial infection^40^, to the best of our knowledge only one phage has been described to target Tad pili in *Caulobacter crescentus*^41^, suggesting that switching from T4a to Tad pili may limit *P. aeruginosa* susceptibility to phage predation.

In this study, we screened for heterogeneously expressed pathways during infection conditions in physiologically-relevant models of the airway epithelium. We identified a number of heterogeneously expressed modules, including Tad pili, and established methods to characterize the physiological state of sorted bacterial subpopulations. Bacteria with high levels of Tad pili co-express QS-related virulence factors and inflict higher levels of tissue damage. In contrast, strains lacking Tad pili have difficulties attaching to the host tissue and lead to slower destruction of the epithelium. Lastly, we identified that a subset of *P. aeruginosa* clinical isolates also expresses Tad pili in a heterogeneous manner. This study highlights the diversity of virulence mechanisms employed by *P. aeruginosa* to damage the host tissue, where different subsets of bacteria may use different strategies to inflict damage.

## Materials and methods

### Bacterial strains and growth conditions

Bacterial strains used in this study can be found in Supplementary Table 9. Bacteria were grown in LB at 37°C for overnights and *in vitro* assays, and in SCFM for day cultures and prior to infection. *E. coli* DH5α λpir was used as cloning strain and *E. coli* SM10 λpir was used for conjugation of *P. aeruginosa*. Antibiotics used were gentamicin (20 μg/mL for *P. aeruginosa* and *E. coli*) and kanamycin (50 μg/mL for *E. coli* and 500 μg/mL for *P. aeruginosa*).

### Mutant construction

*P. aeruginosa* mutants were constructed as previously described using the pEXG2 suicide plasmid. Briefly, 800bp homology arms were amplified by PCR using primers (a list of all primers can be found in Supplementary Table 10) with 20bp stretches that were homologous to the HindIII and XbaI restriction sites of the pEXG2 plasmid. PCR products were inserted into linearized pEGX2 plasmid by Gibson assembly (NEB, E2611L). The resulting vectors were transformed into chemically-competent E. coli DH5α λpir, selected on LB plates with 20 μg/mL gentamicin and verified by colony PCR and Sanger sequencing. The purified plasmids were then transformed into chemically-competent E. coli SM10 λpir and the resulting strains were used for conjugation of P. aeruginosa PAO1. Selection of P. aeruginosa transconjugants was performed on LB with 20 μg/mL gentamicin and 20 μg/mL irgasan. The pEXG2 backbone was then curated by restreaking on LB plates without salt supplemented with 20 μg/mL irgasan and 10% sucrose, which were incubated for 36h at room temperature. Colonies were screened by colony PCR with flanking primers.

pSEVA231-derived plasmids were constructed by insertion of the *mNeongreen* or *mGreenlantern* genes into the HindIII/BamHI restriction sites by Gibson assembly as described above. Sequences of reporters of interest were purchased as eBlocks® (IDT DNA) and cloned upstream from the *mNeongreen* and *mGreenlantern* genes into the EcoRI/SacI restriction sites. The P*_rplM_*-*mCherry2* constitutively expressed fusion was inserted in a convergent orientation using the HindIII/SpeI restriction sites. pPSV35-derived plasmids were constructed by insertion of the genes of interest into the BamHI/EcoRI restriction sites by Gibson assembly and sequence-verified by Sanger sequencing. All plasmids were introduced into P*. aeruginosa* by electroporation of 20-200ng of plasmid into 1 mL of overnight stationary phase cultures washed twice with sterile water using a Bio-Rad GenePulser with the following settings: 25 μF, 400 Ω, 1.8 kV. All steps were performed at room temperature. 1 mL of LB medium was added immediately after the pulse and cells were incubated 1.5-2h at 37°C with shaking before plating on LB plates containing 500 μg/mL kanamycin or 20 μg/mL gentamicin as appropriate.

The pSEVA231-derived plasmids were introduced into clinical isolates of *P. aeruginosa* via conjugation using *E. coli* JKE201 as the donor strain as described^42^. JKE201 was cultured in LB medium supplemented with 1 mM diaminopimelic acid (DAP). For conjugation, concentrated overnight cultures of donor and recipient were mixed at a ∼1:1 ratio on solid media and incubated at 37°C for 4 hours. Mating mixtures were then plated on selective medium (500 µg/mL kanamycin) lacking DAP, to select for clinical isolates that acquired the reporter plasmid and to counterselect against the DAP-auxotrophic donor.

### Flow cytometry

Bacteria carrying fluorescent reporter plasmids were either (i) grown to mid-exponential phase at 37°C for 3h in SCFM or LB, (ii) grown to stationary phase for 24h in SCFM, or (iii) used to infect A549 or the NHBEC model from mid-exponential phase cultures in SCFM. Prior to measurement, SCFM-grown samples were diluted 100x in PBS; A549 samples were lysed with 100 µL 1% Triton X-100 in water; and NHBEC model samples were disrupted by glass bead beating for 30s on a vortex, fixed with 2% PFA for 10 min at RT, washed once in PBS and diluted 10x in PBS. Samples were analysed in a BD Fortessa flow cytometer, with blue laser at different voltages and green laser at 950 volts in low flow rate mode. Data were analysed using FlowJo v10, and the median fluorescence intensity (MFI) was plotted using Prism.

For the reporter screen, the distributions of scaled GFP-intensity values were analysed in R. Briefly, Hartigan’s dip test of unimodality was performed^43^, and skewness, kurtosis and the bimodality coefficient^44^ of the samples were calculated. Promoters with a dip t-test < 0.05 and bimodality coefficient >0.555 were considered as significant (non-normal/bimodal distributions).

### Cell sorting

Fluorescently labelled bacteria were sorted from heterogeneity-inducing conditions such as SCFM or after infection of the NHBEC model. The model was disrupted by glass bead beating for 30s on a vortex in 1 mL PBS. Samples were centrifuged on a tabletop centrifuge at maximum speed at RT, fixed with 2% PFA for 10 min at RT, washed with PBS and sorted with a FACS AriaIII cell sorter equipped with a 70µm nozzle, yielding 50 µL of sample for 50,000 sorted bacteria and 2 mL for 2·10^6^ bacteria. For low input proteomics, bacteria were sorted directly into 5X lysis buffer containing 0.05% DDM (n-dodecyl β-maltoside), 50 mM TCEP (Tris (2-carboxyethyl)phosphine hydrochloride), 500 mM TEAB (triethylammonium bicarbonate) and 75 mM CAA (chloroacetamide) and stored at −80°C until further processing.

### NHBEC-ALI model generation

NHBEC inserts were generated from NHBE cells (Lonza, donor 22TL332331) as previously described^17^. Briefly, NHBE cells were thawed and expanded in PneumaCult™-Ex Plus medium (#05040, STEMCELL Technologies) until confluence on a flask pre-coated for 24h with type I collagen (40 µg/mL in water). PneumaCult™-Ex Plus medium was supplemented with PneumaCult™-Ex Plus 50X supplement, 1000X-diluted hydrocortisone (#7926, STEMCELL Technologies). Next, 33,000 cells/insert were seeded onto 6.5 mm transwell plates (#3470, Corning) and incubated for 2-3 days until confluence with PneumaCult™-Ex Plus medium. The cells were then transferred to an air-liquid interface (ALI) by removal of the medium from the apical side and replacement of the basal medium with PneumaCult™-ALI maintenance medium (supplemented with PneumaCult™-ALI 10X supplement, 200X-diluted hydrocortisone and 0.0004% heparin (#7980, STEMCELL Technologies). The basal medium was exchanged every 2-3 days until the samples were used. From day 28 post-airlift, when the epithelium is fully differentiated, a mucus wash was performed every 7 days by inverted centrifugation at 546g for 5 min at room temperature.

### NHBEC-ALI model infection

Tissues were infected as described^17^. Briefly, *P. aeruginosa* was grown to mid-exponential phase in SCFM and 1000 CFUs in 2 µL SCFM were used to infect the tissues unless otherwise stated. Infected tissues were incubated at 37°C with 5% CO_2_ until samples were fixed and processed for flow cytometry, fluorescence microscopy or proteomics.

### Fluorescence microscopy

At different timepoints post-infection, NHBEC tissue samples were processed as described. Briefly, they were fixed from the apical and basal compartments with 10% formalin for 45 min at RT, washed twice with HBSS (Gibco, #14025092), permeabilized with 0.5% Triton X-100 for 20 min at RT, washed with HBSS and stained for 3h with Hoechst 33342 (1:1000 dilution) and phalloidin-AlexaFluor647 (A22287, ThermoFisher scientific, 1:250 dilution). For staining of the ciliary basal membrane, α-ARL13B rabbit antibodies were used (177111AP150UL, Proteintech, 1:400 dilution) for 3h at RT, followed by 3.5h incubation with goat anti-rabbit Alexa Fluor Plus 488 antibodies (A32723, Life Technologies, 1:500 dilution). Samples were washed four times in HBSS with 0.05% sodium azide and embedded in ProLong Glass AntiFade mountant (P36984, Thermofisher scientific). Samples were curated in the dark for >48h. Samples were imaged on a Nikon ECLIPSE Ti2 microscope with the Crestoptics X-Light V3 spinning disk confocal system with a 60x oil objective (NA 1.42).

### Image analysis

To improve segmentation, self-supervised denoising was performed on RAW image stacks using the N2V2 architecture^45^ implemented via the CAREamics framework^46^. Independent models were trained for each channel. The networks were configured and trained using the following parameters: patch size: 32×256×256 pixels (ZYX), training duration: 100 epochs, validation: 15% of the input data was randomly reserved for validation. The denoising inference used a tile size of 32×256×256 pixels and a tile overlap of 8×64×64 pixels (ZYX).

Nuclei and cells were segmented using the Cellpose-SAM model^47^. The specific hyperparameters for each target were selected as follows (the rest was left as default). For nuclei, 50.00 px diameter, −3.742 flow threshold, 2.43 anisotropy factor and 1.74 3D smooth threshold.

For Cells, 76.27 px diameter, −3.404 flow threshold, 2.786 anisotropy factor, 0.69 3D smooth threshold. Bacteria were segmented using StarDist3D^48^. The StardistOPP model 7 was used with its default probability (0. 45) and non-maximum suppression thresholds (0.3) settings^49^.

To analyse the spatial distribution and structural orientation of the bacterial masks, we used the scikit-image library to find the boundaries of each bacterium, turning the masks into a 3D point cloud. The longest axis of the mask was determined using singular value decomposition. Next, for each bacterium using a k-dimensional tree analysis, we determined how many bacteria were within a 10 µm radius. Finally, the exact distance from the outer shell of a target bacterium to the closest point of its nearest neighbour was calculated as well.

To measure the distances of the bacteria to the host cells, the surfaces of both the cells and the bacteria were extracted using the find_boundaries function from the scikit-image library^50^, turning the masks into 3D boundary point clouds. Using a k-dimensional lookup tree (scipy.spatial.KDTree^51^) the minimum, maximum, and average distance from the bacterial surface to the nearest cellular surface was determined.

The scikit-image library^50^ was used to calculate the 3D centre of mass and total volume for each bacterium. To track shape distortion along the vertical axis, a Z-elongation ratio was calculated by dividing the height of each bacterium by its width. The average brightness of the pixels inside each bacterial mask was calculated using scikit-image and multiplied by the total pixel count to determine the total absolute amount of signal.

To quantify ciliary signals, 3D Euclidean distance maps and Voronoi tessellation were constructed from the segmented bacterial mask using scipy.ndimage.distance_transform_edt, calibrated to physical voxel dimensions. To prevent double-counting cilia signals, each extracellular voxel was mapped to the identity of its closest bacterium using scipy.ndimage.distance_transform_edt, splitting the volume at the midpoint boundary between adjacent cells so only the signal in the neighbouring environment of each bacterium was used for the calculations. For each bacterium, the halo was isolated by selecting voxels within its assigned Voronoi domain having an Euclidean distance between 0 and 3 μm. The mean cilia intensity was determined by dividing the halo intensity sum by its total voxel count. Bacteria whose centroids or bounding limits lay within 3 µm of the FOV border were flagged and excluded from analysis.

### Proteomics sample preparation (STRAP)

At 24 hpi, tissues were cut out of the transwell insert and frozen at −80 °C until further processing. Next, samples were washed once with PBS. From here on, bacteria-only samples that were quantified with the Orbitrap platform were frozen and processed in the same way. Samples were resuspended in 100 µL lysis buffer (5% SDS, 100 mM triethylammonium bicarbonate (TEAB) and 10 mM tris(2-carboxyethyl)phosphine (TCEP)) and boiled for 10 min at 95 °C. The supernatant was transferred to a 96-well U-bottom plate and lysed for 20 min in a PIXUL Multi-Sample sonicator (Active Motif) at 15 °C. Per sample, 10 µg protein were alkylated using 15 mM iodoacetamide at 25° C in the dark for 30 min. The entire protein lysate was captured, digested (trypsin 1/50, w/w; Promega), and desalted using STRAP cartridges (Protifi) following the manufacturer’s instructions. Samples were dried under vacuum and stored at −20 °C until further use.

### Proteomics sample preparation (low input SP3)

At different timepoints post-infection, bacterial cultures or infected tissues were fixed with 2% PFA for 10 min at RT and washed twice with PBS. After that, bacterial samples were diluted to different amounts and lysed with lysis buffer containing 5% SDS, 100 mM TEAB and 10 mM TCEP, similarly like before. Proteins were extracted, reduced and alkylated as described above for s-trap and purified and digested with the SP3 approach^52^ using a Fluent liquid handling platform (Tecan Group Ltd., Männedorf, Switzerland). In brief, Speed BeadsTM (#45152105050250 and #65152105050250, GE Healthcare) were mixed 1:1, rinsed with water and diluted to the 8 μg/µL stock solution. Samples were adjusted to the final volume of 90 µL and 10 µL of the beads stock solution was added to them. Proteins were bound to the beads by addition of 100 µL of 100% acetonitrile (ACN) to the samples, which were then incubated for 8 min at RT with a gentle agitation (200 rpm). After, samples were placed on a magnetic rack and incubated for 5 minutes. Supernatants were removed and discarded. The beads were washed twice with 160 µL of 70 % (v/v) ethanol and once with 160 of 100% ACN. Samples were placed off the magnetic rack and 50 µL of digestion mix (10 ng/µL of trypsin in 50 mM TEAB, 0.01% n-dodecyl-ß-maltoside (DDM)) was added to them. Digestion was allowed to proceed for 12h at 37°C. After digestion samples were placed back on the magnet and incubated for 5 minutes. Supernatants containing peptides were collected and dried under vacuum.

### Proteomics sample preparation (DDM protocol)

Bacterial samples were sorted into 5X lysis buffer containing 0.05% DDM, 50 mM TCEP and 500 mM TEAB in a 96 well low binding PCR plate (Eppendorf) and frozen at −80 °C until further processing. Then, they were processed according to a recently published protocol for very low cell numbers^53^. The samples were heated to 95°C (lid temperature 110°C) for 1 hour in a thermal cycler (BioRad, S1000). After cooling, 8.5 μL of digestion buffer (60% ACN, 100 mM TEAB in water) was added and the samples heated to 75°C for 30 minutes. Then, 1.4 μL of trypsin solution (100 ng/μL trypsin in 50 mM TEAB) were added to the cooled samples and the protein digested for 2 hours at 37°C. After adding 1 μL of 5% trifluoracetic acid (TFA), the samples were dried in a Speedvac. Subsequently, the peptides were cleaned by solid phase extraction using EvoTip Pure stage tips (EVOSEP, EV2020). The dried peptides were resuspended in 10 μL 0.1% TFA using sonication for 5 minutes in a waterbath. The sample were loaded onto EvoTips by centrifugation following the vendor’s protocol. Peptides were eluted with 20 ul elution buffer (40% ACN, 0.1% formic acid (FA)) directly into a new low binding PCR 96-well plate, dried under vacuum and stored at −20 °C until further use.

### Proteomics sample preparation (DDM protocol with alkylation and overnight digestion)

Same as above with the following changes according to^54^: 75 mM CAA was added to the 5X lysis buffer and proteins were digested overnight.

### LC-MS analysis (Orbitrap)

Dried peptides were resuspended in 0.1% aqueous formic acid and subjected to LC– MS/MS analysis using an Orbitrap Eclipse Tribrid Mass Spectrometer fitted with an Ultimate 3000 nano-LC (both Thermo Fisher Scientific) and a custom-made column heater set to 60°C using block randomization. Peptides were resolved using a RP-HPLC column (75μm × 30cm) packed in-house with C18 resin (ReproSil-Pur C18–AQ, 1.9 μm resin; Dr. Maisch GmbH) at a flow rate of 0.3 μLmin-1. The following gradient was used for peptide separation: from 2% B to 12% B over 5 min to 30% B over 40 min to 50 % B over 15 min to 95% B over 2 min followed by 11 min at 95% B. Buffer A was 0.1% formic acid in water and buffer B was 80% acetonitrile, 0.1% formic acid in water.

The mass spectrometer was operated in DIA mode with a cycle time of 3 seconds. MS1 scans were acquired in the Orbitrap in centroid mode at a resolution of 60,000 FWHM (at 200 m/z), a scan range from 350 to 1200 m/z, normalized AGC target set to 250 % and maximum ion injection time mode set to 50 ms. MS2 scans were acquired in the Orbitrap in centroid mode at a resolution of 15,000 FWHM (at 200 m/z), precursor mass range of 400 to 900, quadrupole isolation window of 12 m/z with 1 m/z window overlap, a defined first mass of 120 m/z, normalized AGC target set to 1000 % and a maximum injection time of 22 ms. Peptides were fragmented by HCD (Higher-energy collisional dissociation) with collision energy set to 33 % and one microscan was acquired for each spectrum.

### LC-MS analysis (timsTOF)

Peptides were resuspended in 5 ul 0.1% FA and the whole sample injected for LC–MS/MS analysis using a timsTOF Ultra 2 Mass Spectrometer (Bruker) fitted with a Vanquish Neo HPLC (Thermo Fisher Scientific). The column heater was set to 60°C. Peptides were resolved using a RP-HPLC column (100μm × 30cm) packed in-house with C18 resin (ReproSil Saphir 100 C18, 1.5 μm resin; Dr. Maisch GmbH) at a flow rate of 0.4 μL min-1 at 60°C. A linear gradient of buffer B (80% ACN, 0.1% formic acid in water) ranging from 2% to 25% over 25 minutes and from 25% to 35% over 5 minutes was used for peptide separation. Buffer A was 0.1% FA in water. The mass spectrometer was operated in positive DIA-PASEF mode. MS spectra were acquired from 100 to 1700 m/z. For MS/MS, we used eight MS/MS ramps and 24 MS/MS windows covering a mass-to-charge ratio range of 400–1000 m/z. This resulted in 25 Th wide MS windows and an ion mobility range from 0.64 to 1.37 V -s/cm². The mass spectrometer was operated in normal sensitivity mode, expected cycle time was 0.95 seconds, accumulation and ramp time was set to 100 ms. The following source settings were applied: 1600V Capillary Voltage and 3 l/min Dry Gas flow at 200°C. The collision energy was linearly ramped between 20 eV at 0.6 V -s/cm² and 59 eV at 1.6 V -s/cm².

### Proteomics data analysis

The acquired raw files were searched using SpectroNaut (Biognosys, Schlieren, Switzerland, v19.0, default settings) against a *P. aeruginosa* database (consisting of 5,564 protein sequences downloaded from Uniprot on 2021/07/02) and 393 commonly observed contaminants using the following search criteria: full tryptic specificity was required (cleavage after lysine or arginine residues, unless followed by proline); 3 missed cleavages were allowed; carbamidomethylation (C) was set as fixed modification; oxidation (M), N-acetlyation (N-term) were applied as variable modifications. The raw quantitative data was further statically analyzed using ProteoFlux (v1.8.7)^55^ and Prism (v10.3.1, GraphPad). ProteoFlux is an open-source and reproducible workflow for quantitative proteomics data analysis. Prior to quantification, precursor intensities were filtered based on run-level posterior error probability (PEP) and q-value thresholds (≤ 0.01), and very low-intensity signals (below a threshold of 2) were treated as left-censored measurements. Precursor-level intensities exported from Spectronaut were aggregated to peptide-level quantities and subsequently summarized to protein abundances using a simple summarization-based approach, followed by median-based sample normalization and statistical analysis using linear models with empirical Bayes moderation.

Enrichment analyses were performed as follows. Gene classes were obtained from KEGG and GO Biological Process databases and pruned by Jaccard’s similarity, where pathways containing >50% shared genes are aggregated. Next, resulting GMT files were used to perform enrichment analysis of the proteomics output. Bacterial samples were analyzed using the fgsea package^56^ that accurately detects mild changes of full regulons, using the full list of detected bacterial proteins and their log fold changes as input. For human samples, hypergeometric enrichment analysis was performed instead^57^, which is more sensitive to targeted proteome changes rather than coordinated transcriptional responses, using hits with a q-value < 0.05 as significantly dysregulated.

### Visualization and processing of RNA-seq data from the BACTOME transcriptomic dataset

A matrix containing raw read counts for all mapped genes across 414 clinical isolates was downloaded from the BACTOME transcriptomic dataset (GEO Accession number 123544). All normalization and processing steps were performed within the R statistical environment (v 4.6.1) using the edgeR package (Version 4.10.1)^58^ as described previously^59^. Briefly, raw reads were converted to counts per million (CPM) and only genes with ≥1 CPM in at least 21 isolates were retained. The resulting matrix of CPM values was used for normalization using Trimmed Mean of M-values (TMM) method.

Normalized log2-CPM values were scaled by converting to Z-scores calculated per gene. For visualizing expression differences for each gene across isolates, heatmaps were constructed using R packages – heatmaply (Version 1.6.0)^60^ and pheatmap (Version 1.0.13)^61^. Hierarchical clustering for the heatmaps utilized Euclidian distances and complete linkage. The color gradient was weighed to place white at zero, spanning from green (below-mean expression) to magenta (above-mean expression).

Co-expression analysis was performed in R by computing Spearman rank correlation coefficients between the normalized log2-CPM value of each gene and that of the gene of interest across all clinical isolates. This was followed by Benjamini-Hochberg false discovery rate correction using the p.adjust() function of the base R stats package. Results were visualized as a volcano plot of Spearman rho against −log10 FDR-adjusted p-value using the ggplot2 package (v4.0.3)^62^.

### Statistics

When only two means were compared, we performed unpaired t-tests. When multiple means were compared, we used two-way ANOVA with Sidak’s multiple comparison test or the Kruskal-Wallis test with Dunn’s multiple comparisons correction, using GraphPad Prism version 10.2.0. Unless otherwise stated, data are represented as mean ± standard deviation (SD). *p-value<0.05, **p-value<0.005, ***p-value<0.0005, ****p-value<0.00005, ns – p-value>0.05.

## Data availability

All mass spectrometry proteomics data associated with this manuscript have been deposited to the ProteomicsXchange consortium via MassIVE (https://massive.ucsd.edu) with the accession number MSV000103361/PXD084749.

## Acknowledgements

This work was supported by the National Center of Competence in Research AntiResist funded by the Swiss National Science Foundation (51NF40_180541) to MB and UJ. VS was supported by Biozentrum PhD fellowships. We thank Benoît Laventie and Leoni Swart for discussions and advice on the airway model and Sascha Siegfried for technical help. We thank Laurent Guerard for advice on the image analysis pipeline and Susanne Häussler for the 414 clinical isolate collection. We thank Anamaria Derek for advice on ciliary staining. We thank Stella Stefanova and Svitlana Malysheva of the Biozentrum Flow Cytometry Facility for help with flow cytometry and Matthias Ostermaier from the Proteomics Core Facility for experimental help.

## Author Contributions

Conceptualization: ATA, MB; Methodology: ATA, UL, RF, AS; Software: ATA, VS, EB, OB, MB; Validation: ATA, VS; Formal analysis: ATA, VS, MAV, AS; Investigation: ATA, VS, AS; Resources: UJ, AS, MB; Data curation: ATA; Writing – original draft: ATA; Writing – review & editing: ATA, MB; Visualization: ATA, VS, EB; Supervision: ATA, OB, AS, MB; Project administration: ATA, MB; Funding acquisition: MB, UJ.

