## Supplementary Figures for "Subpopulations of *Pseudomonas aeruginosa* inflict distinct tissue damage during infection"

### Supplementary Figure 1

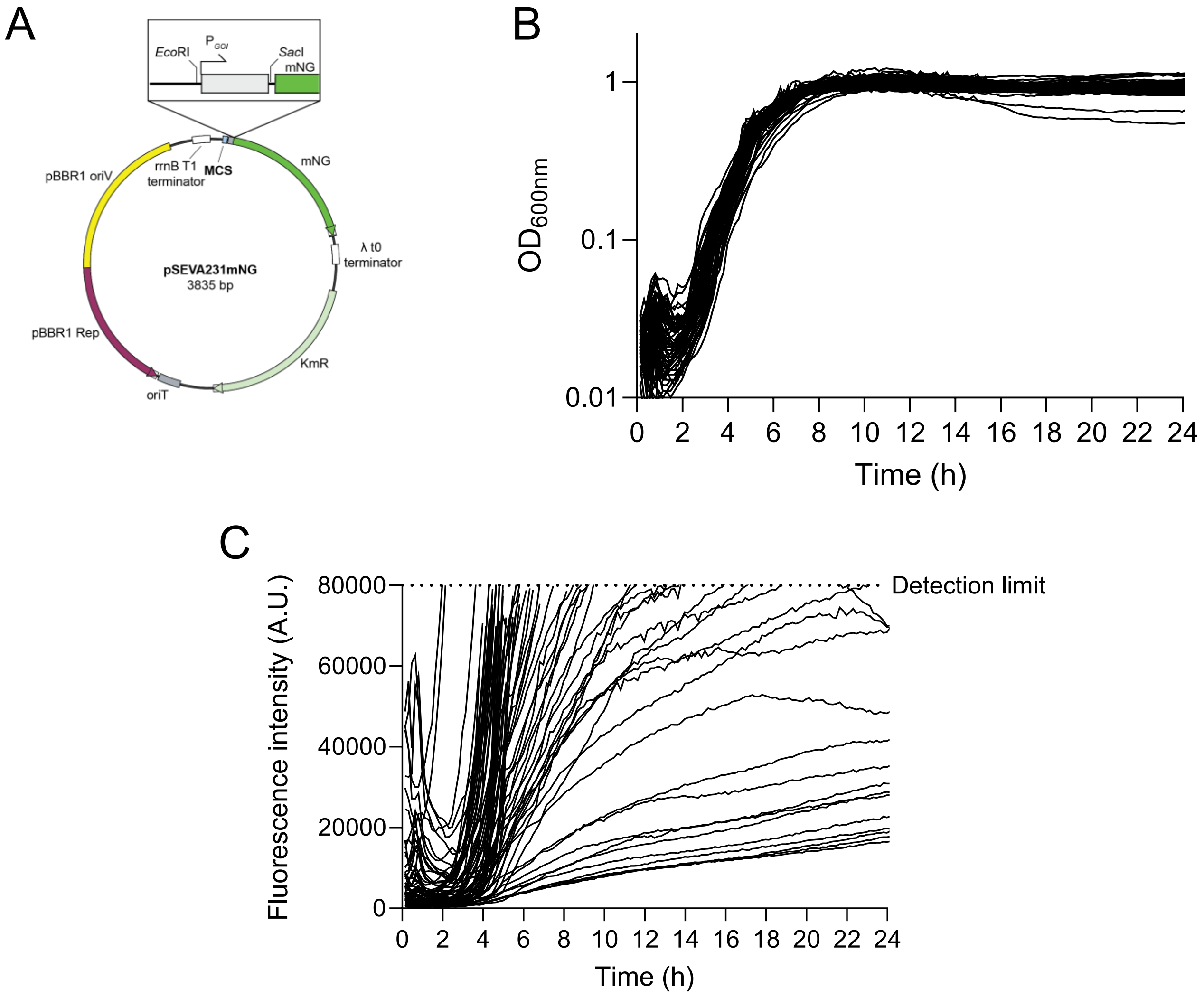

### Supplementary Figure 2

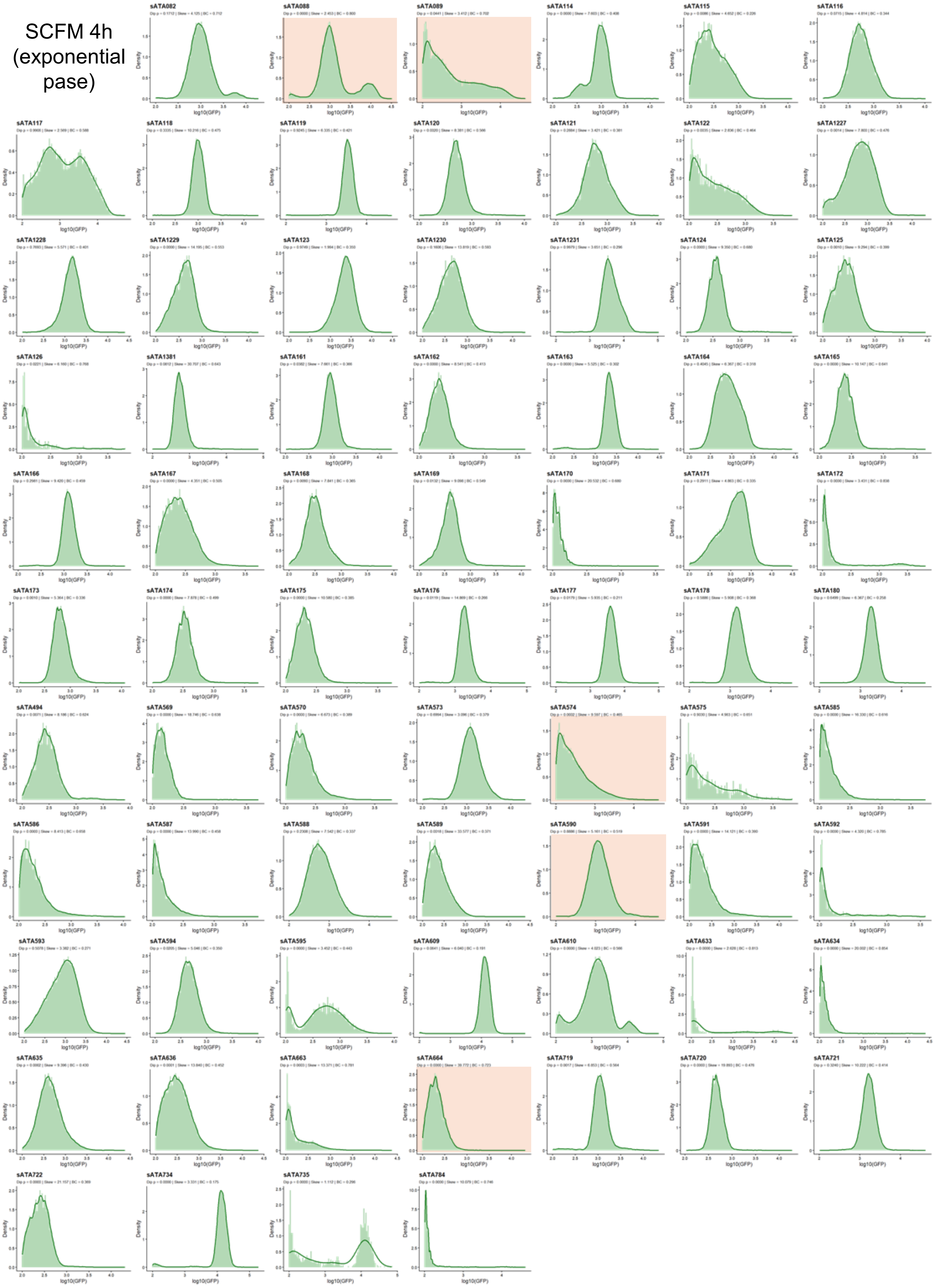

### Supplementary Figure 3

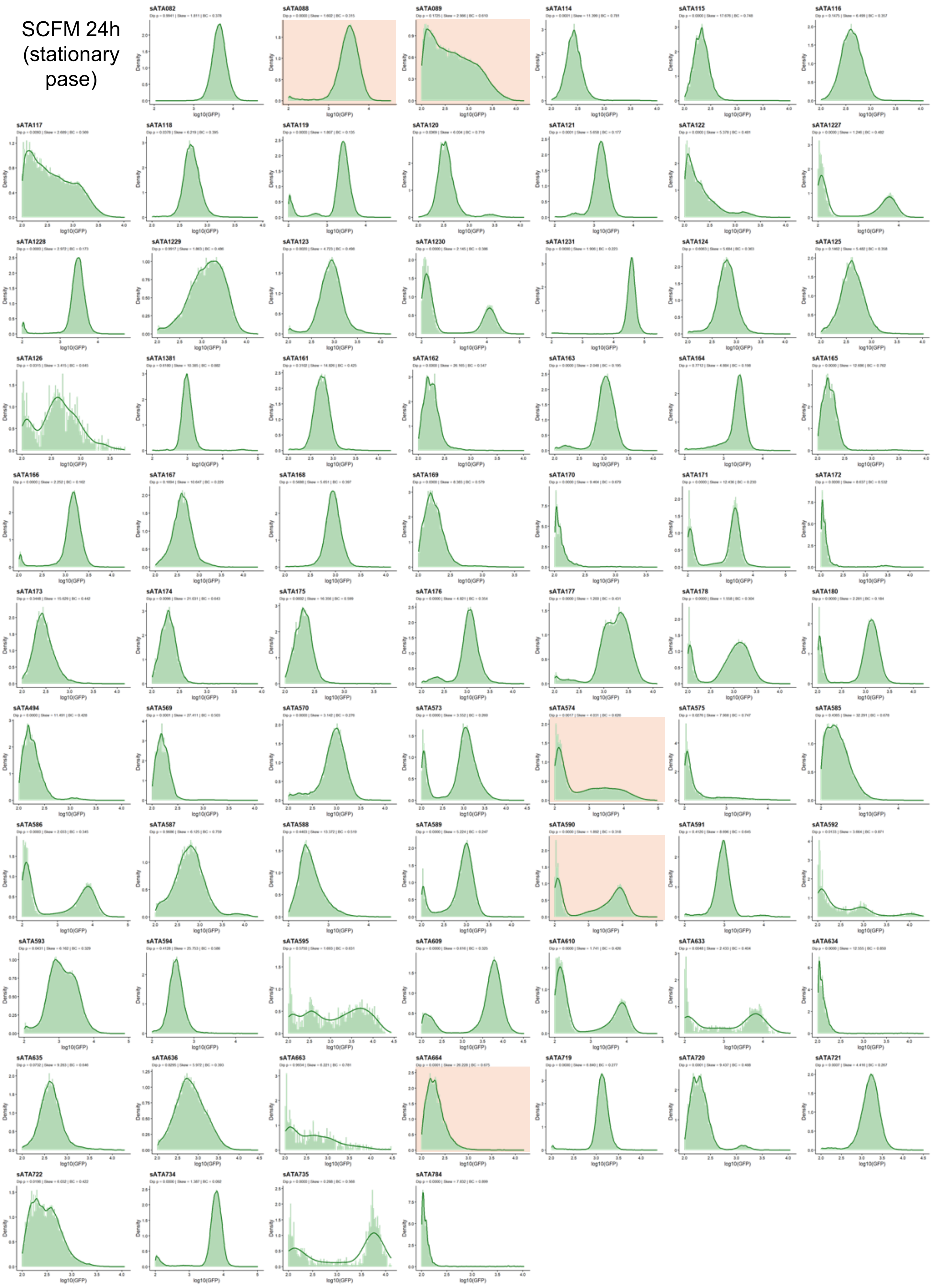

### Supplementary Figure 4

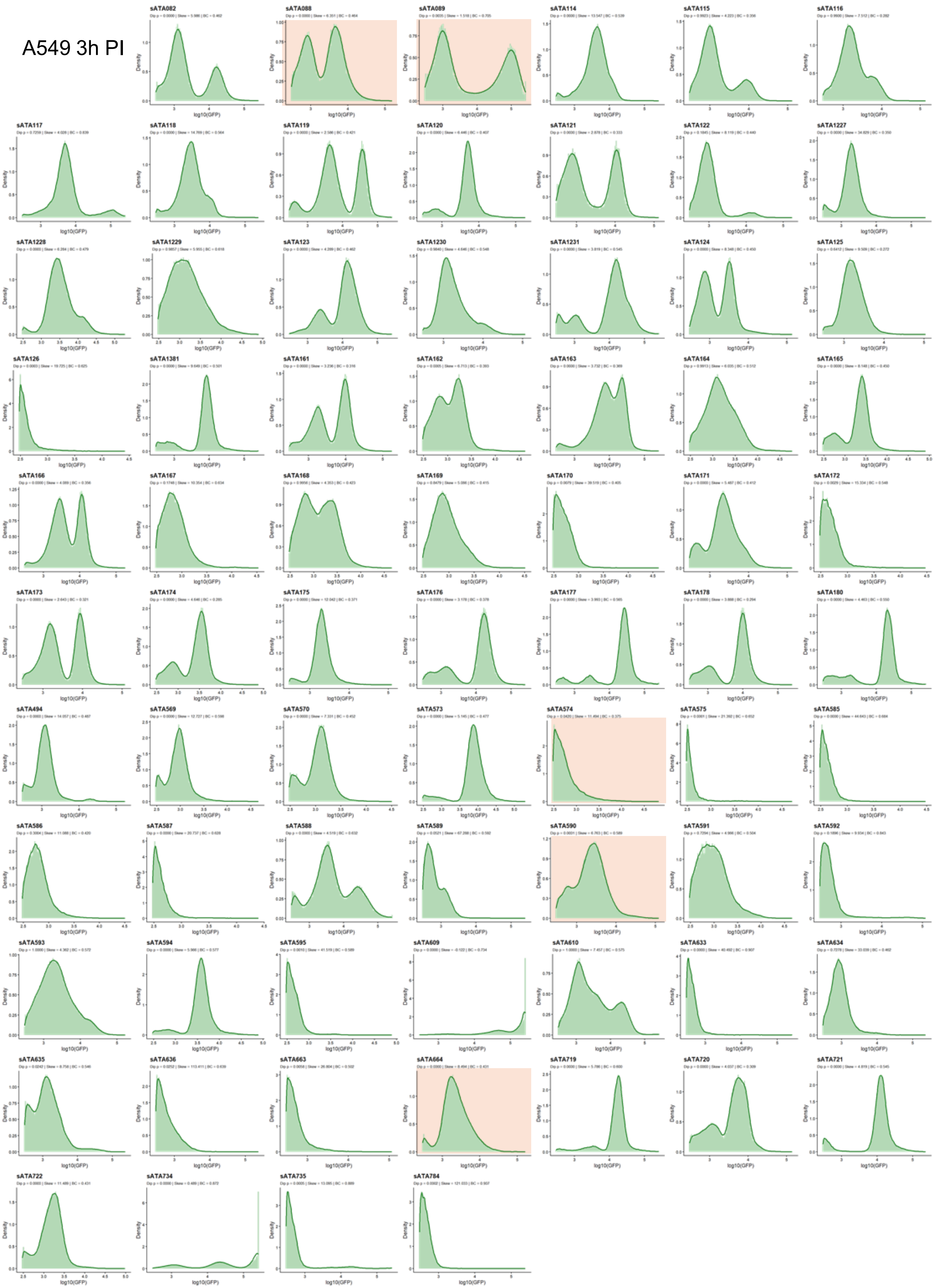

### Supplementary Figure 5

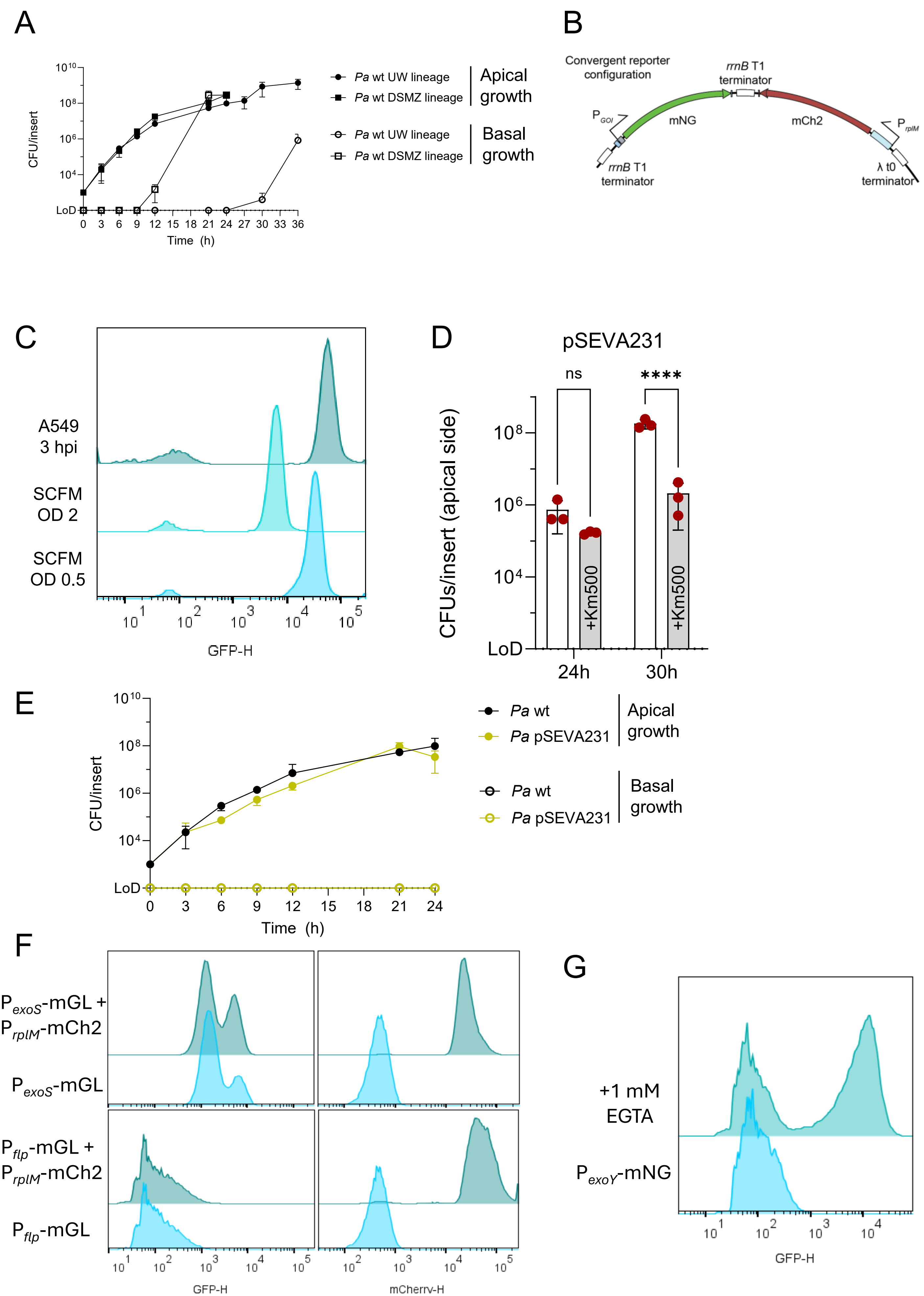

Supplementary Figure 6

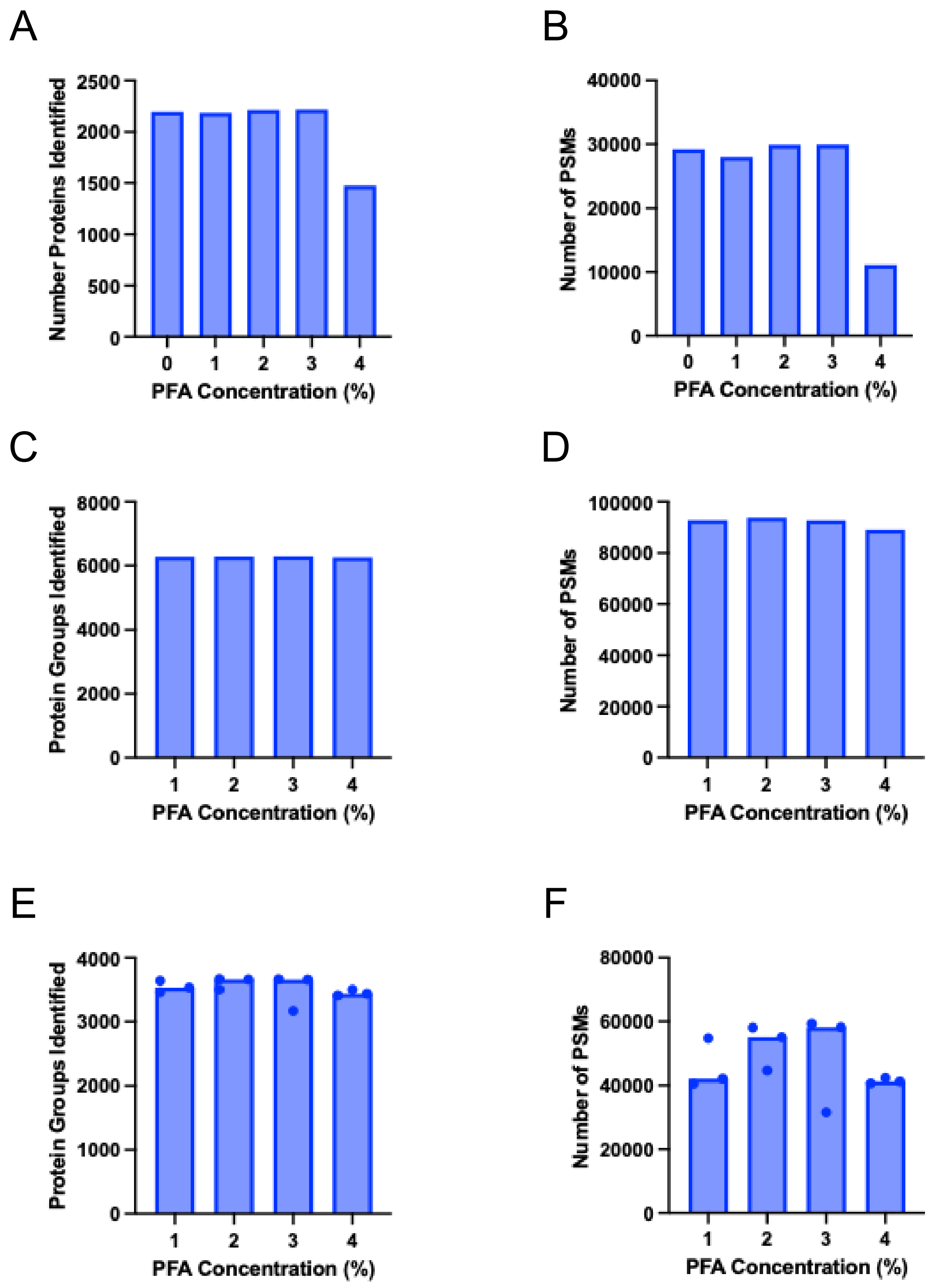

Supplementary Figure 7

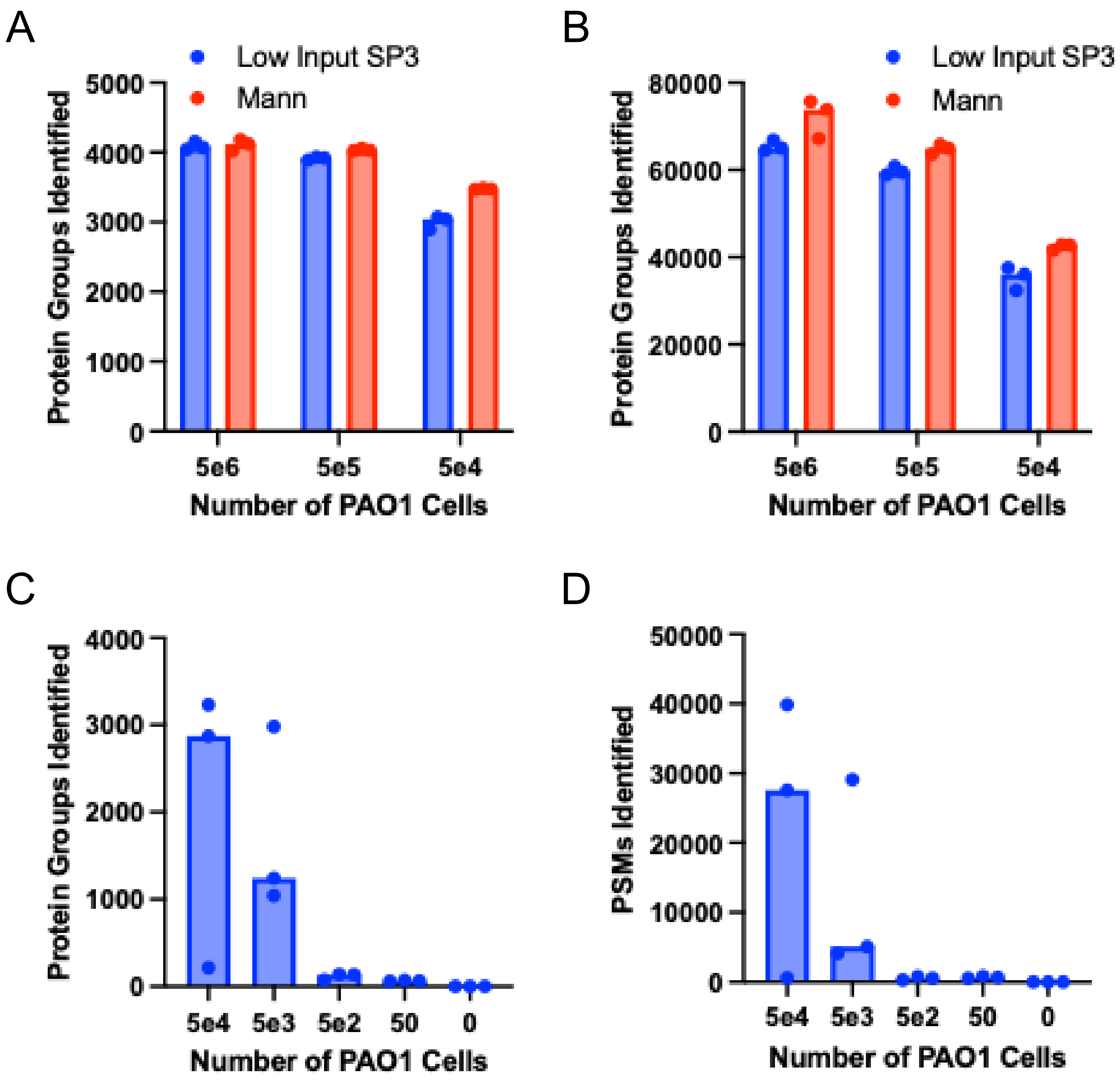

Supplementary Figure 8

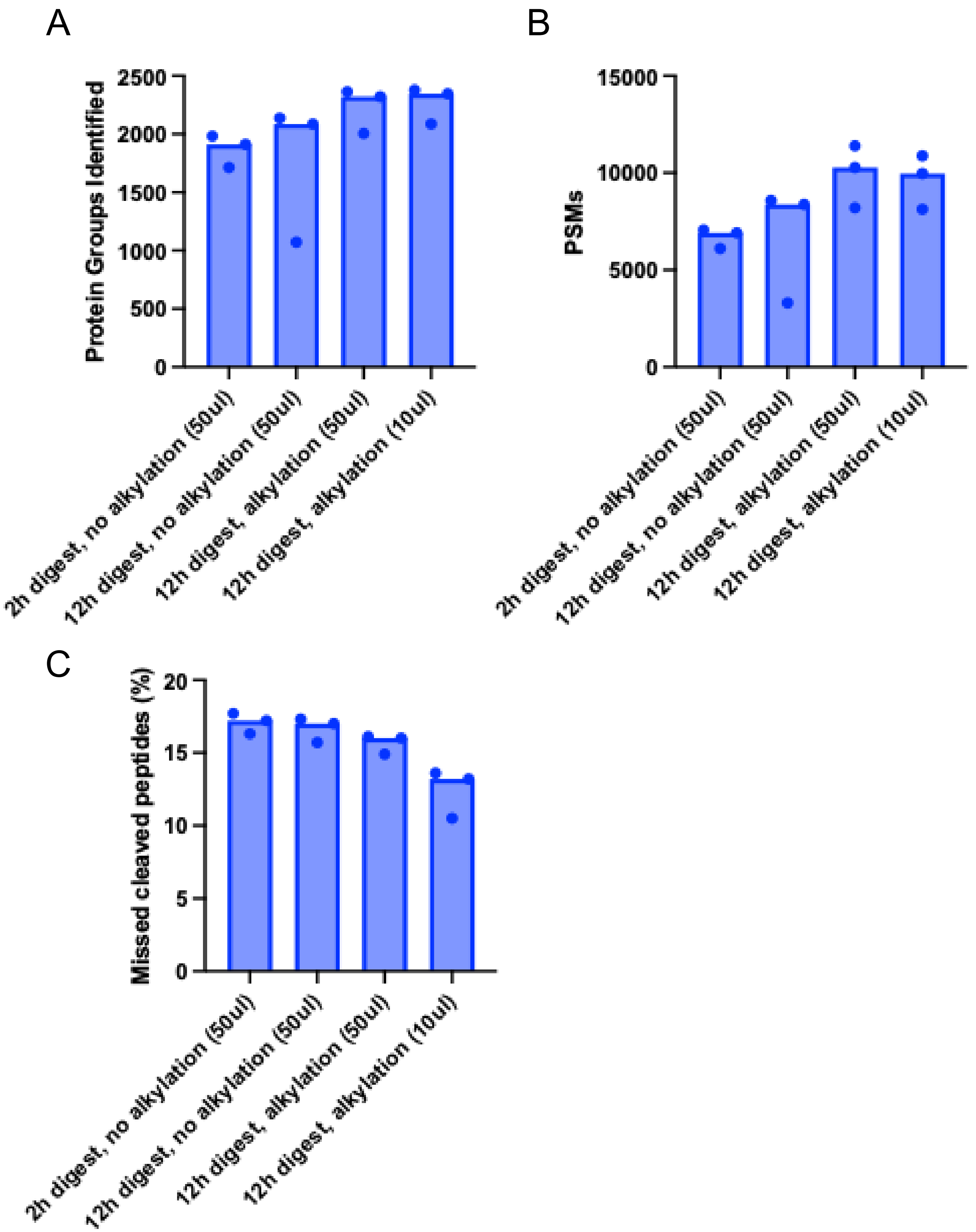

Supplementary Figure 9

A

| Experiment | Number of Bacteria per Sample | Sample Preparation Method* | LC-MS Platform |
|---|---|---|---|
| 1 | 2E6 | s-trap | Orbitrap |
| 2 | 1E8 | s-trap | Orbitrap |
| 3 | 5E6 | Low Input SP3 | timsTOF Ultra 2 |
| 4 | 5E4 | Low Input SP3 | timsTOF Ultra 2 |
| 5 | 5E4 | DDM, 2h digestion | timsTOF Ultra 2 |
| 6 | 5E4 | DDM, alkylation, 12h digestion | timsTOF Ultra 2 |

B

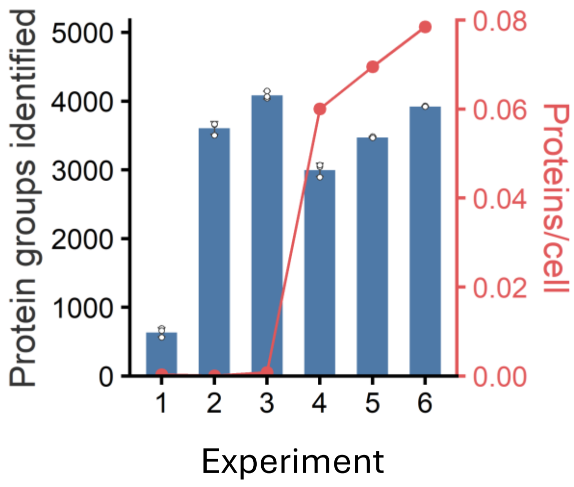


C

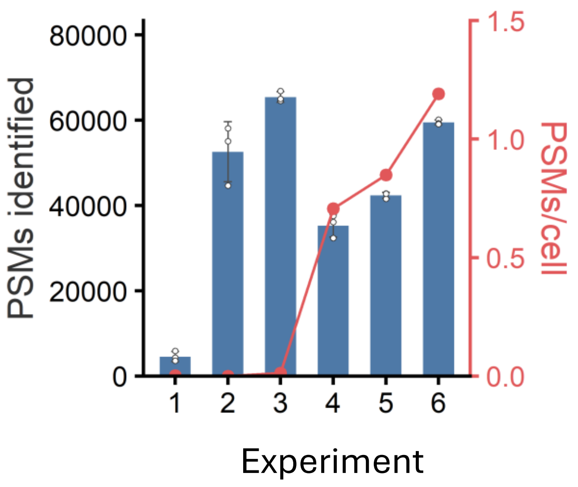


* See method section for details

### Supplementary Figure 10

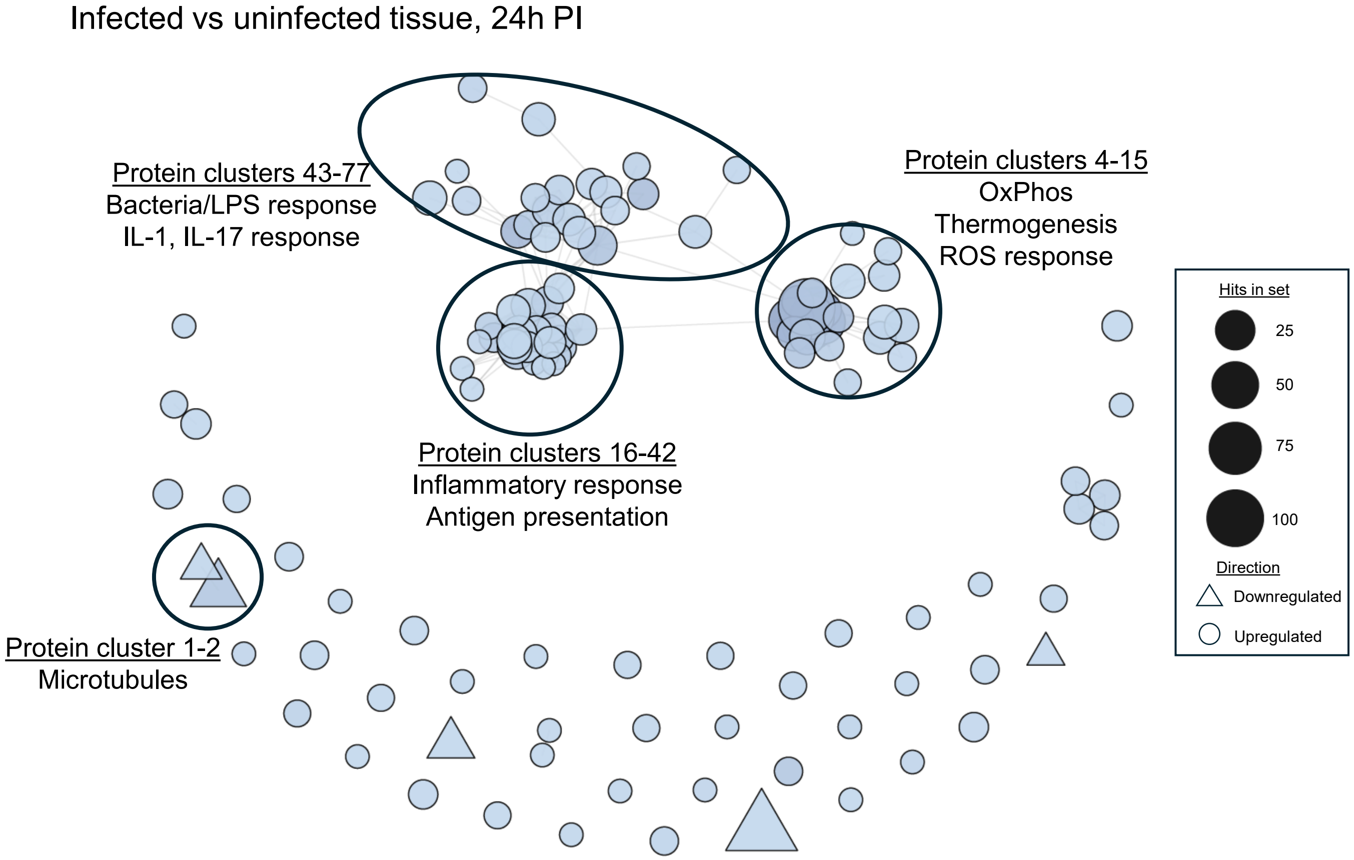

### Supplementary Figure 11

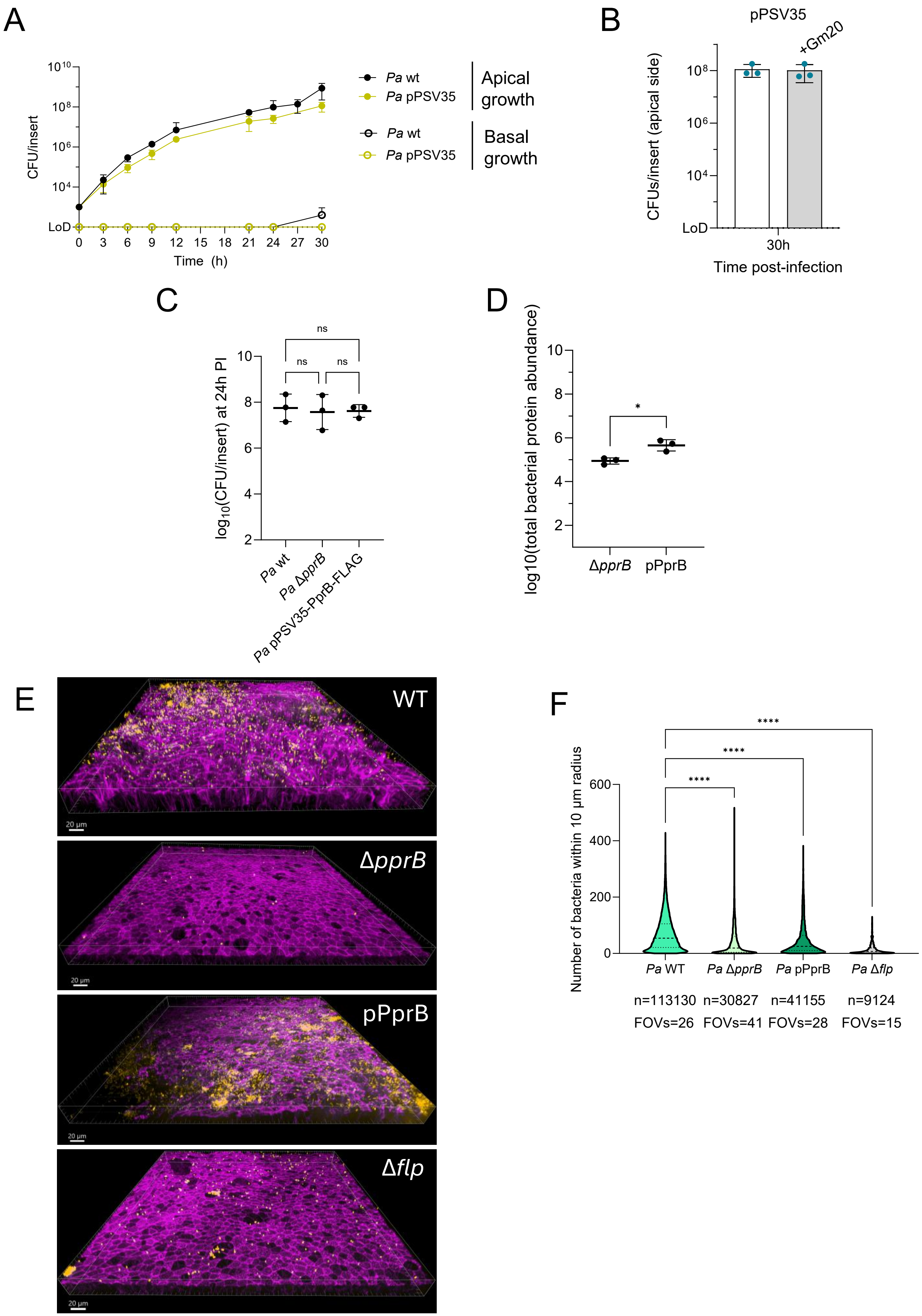

### Supplementary Figure 12

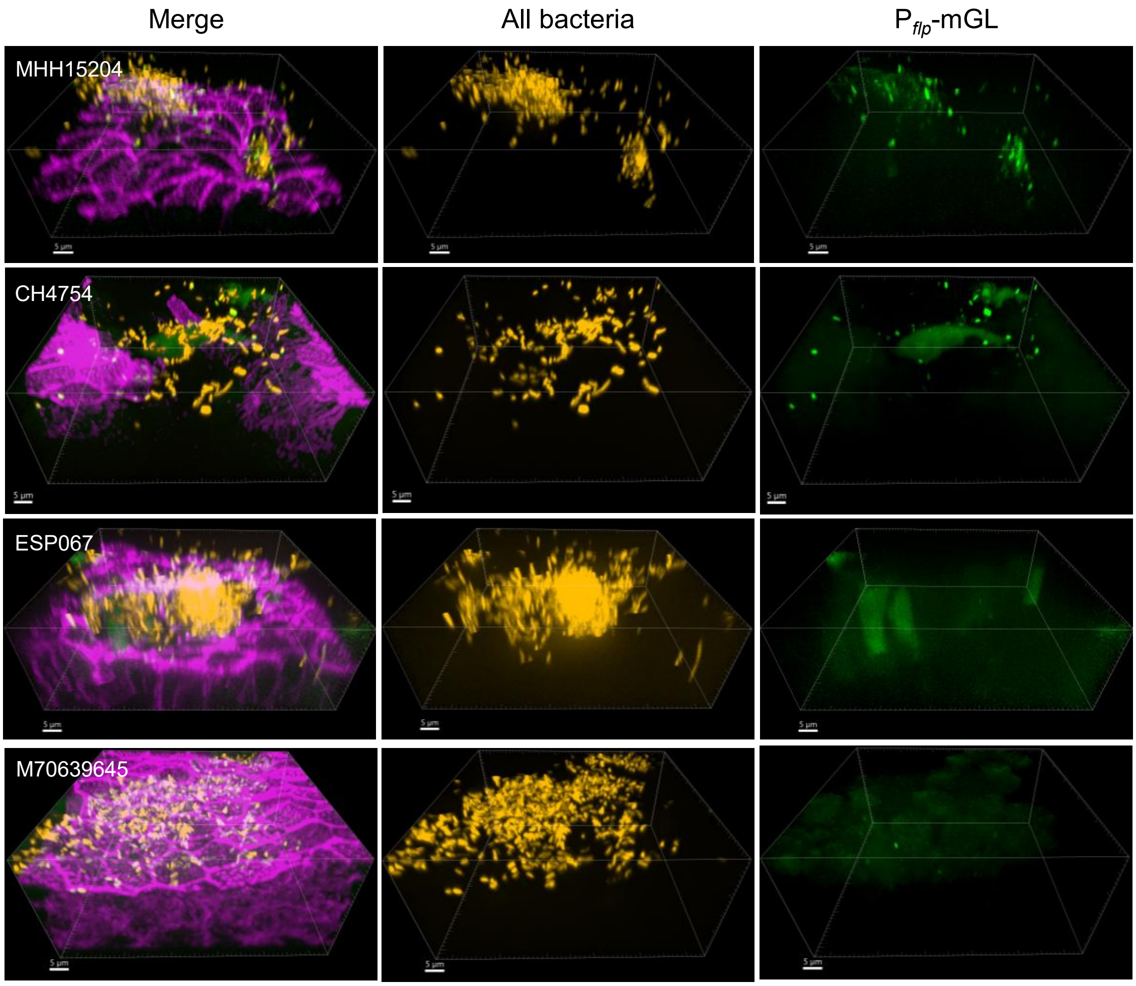
