## Supplementary File 1 for "Subpopulations of *Pseudomonas aeruginosa* inflict distinct tissue damage during infection"

**Supplementary File 1. Details on the optimization of the low input proteomics strategy.**

We compared the sensitivities of the Orbitrap and timsTOF Ultra 2 liquid chromatography mass spectrometry (LC-MS) platforms with different amounts of input bacteria. In addition, we assessed the impact of different fixative concentrations (Supplementary Figure 6), different sample preparation protocols (Supplementary Figure 7) and different protein digestion durations and cysteine alkylation (Supplementary Figure 8) on the number of proteins identified, Peptide-Spectrum Matches (PSMs) and protein groups identified (Supplementary Figures 6-8). A summary of the sensitivity and proteome coverage improvements achieved with the most relevant optimization steps is in Supplementary Figure 9. Using the optimized protocol, we could identify >3,900 protein groups and 1.2 PSMs/cell using as little as 50,000 sorted *P. aeruginosa* cells, which approximately represents a 2,200-fold improvement compared to Orbitrap-based traditional proteomics approaches. Therefore, our method allows the accurate quantification of the proteome of down to 50,000 bacterial cells, enabling the coupling of FACS-based methods and proteomics while bypassing the need for sample concentration.
