## Supplementary Table 9 for "Subpopulations of *Pseudomonas aeruginosa* inflict distinct tissue damage during infection"

**Supplementary Table 2. Strains used in this study.**

| Strain number | Strain genotype | Plasmid | Resistance | Reference |
|---|---|---|---|---|
| *E. coli* | | | | |
| sATA016 | DH5α λpir | | | [1] |
| sATA380 | SM10 λpir | | | [2] |
| sATA067 | CC118 | pSEVA231 | Km50 | [3] |
| sATA069 | DH5α λpir | pSEVA231-mNG | Km50 | This study |
| sATA221 | DH5α λpir | pPSV35 | Gm20 | [4] |
| sATA080 | DH5α λpir | pSEVA231-$P_{algU}$-mNG | Km50 | This study |
| sATA083 | DH5α λpir | pSEVA231-$P_{hpd}$-mNG | Km50 | This study |
| sATA084 | DH5α λpir | pSEVA231-$P_{norC}$-mNG | Km50 | This study |
| sATA100 | DH5α λpir | pSEVA231-$P_{fabD}$-mNG | Km50 | This study |
| sATA085 | DH5α λpir | pSEVA231-$P_{fhp}$-mNG | Km50 | This study |
| sATA102 | DH5α λpir | pSEVA231-$P_{flhA}$-mNG | Km50 | This study |
| sATA092 | DH5α λpir | pSEVA231-$P_{nirS}$-mNG | Km50 | This study |
| sATA094 | DH5α λpir | pSEVA231-$P_{nuoA}$-mNG | Km50 | This study |
| sATA095 | DH5α λpir | pSEVA231-$P_{pa0797}$-mNG | Km50 | This study |
| sATA414 | DH5α λpir | pSEVA231-$P_{recN}$-mNG | Km50 | This study |
| sATA099 | DH5α λpir | pSEVA231-$P_{fdnG}$-mNG | Km50 | This study |
| sATA097 | DH5α λpir | pSEVA231-$P_{narK1}$-mNG | Km50 | This study |
| sATA101 | DH5α λpir | pSEVA231-$P_{pilA}$-mNG | Km50 | This study |
| sATA098 | DH5α λpir | pSEVA231-$P_{aceA}$-mNG | Km50 | This study |
| sATA093 | DH5α λpir | pSEVA231-$P_{lldA}$-mNG | Km50 | This study |
| sATA113 | DH5α λpir | pSEVA231-$P_{oprO}$-mNG | Km50 | This study |
| sATA110 | DH5α λpir | pSEVA231-$P_{cifR}$-mNG | Km50 | This study |
| sATA145 | DH5α λpir | pSEVA231-$P_{pa2469}$-mNG | Km50 | This study |
| sATA153 | DH5α λpir | pSEVA231-$P_{aspS}$-mNG | Km50 | This study |
| sATA138 | DH5α λpir | pSEVA231-$P_{pchR}$-mNG | Km50 | This study |
| sATA154 | DH5α λpir | pSEVA231-$P_{cspA}$-mNG | Km50 | This study |
| sATA108 | DH5α λpir | pSEVA231-$P_{pa3754}$-mNG | Km50 | This study |
| sATA107 | DH5α λpir | pSEVA231-$P_{bkdA1}$-mNG | Km50 | This study |
| sATA146 | DH5α λpir | pSEVA231-$P_{fiuI}$-mNG | Km50 | This study |
| sATA137 | DH5α λpir | pSEVA231-$P_{phuR}$-mNG | Km50 | This study |
| sATA142 | DH5α λpir | pSEVA231-$P_{ampC}$-mNG | Km50 | This study |
| sATA105 | DH5α λpir | pSEVA231-$P_{fagA}$-mNG | Km50 | This study |
| sATA140 | DH5α λpir | pSEVA231-$P_{pa5539}$-mNG | Km50 | This study |
| sATA143 | DH5α λpir | pSEVA231-$P_{oprD}$-mNG | Km50 | This study |
| sATA147 | DH5α λpir | pSEVA231-$P_{mexA}$-mNG | Km50 | This study |
| sATA148 | DH5α λpir | pSEVA231-$P_{muxA}$-mNG | Km50 | This study |
| sATA141 | DH5α λpir | pSEVA231-$P_{lexA}$-mNG | Km50 | This study |
| sATA150 | DH5α λpir | pSEVA231-$P_{groES}$-mNG | Km50 | This study |
| sATA149 | DH5α λpir | pSEVA231-$P_{ibpA}$-mNG | Km50 | This study |
| sATA144 | DH5α λpir | pSEVA231-$P_{betI}$-mNG | Km50 | This study |
| sATA139 | DH5α λpir | pSEVA231-$P_{pchE}$-mNG | Km50 | This study |
| sATA519 | DH5α λpir | pSEVA231-$P_{pa0830}$-mNG | Km50 | This study |
| sATA520 | DH5α λpir | pSEVA231-$P_{coxB}$-mNG | Km50 | This study |
| sATA523 | DH5α λpir | pSEVA231-$P_{tssA1}$-mNG | Km50 | This study |
| sATA524 | DH5α λpir | pSEVA231-$P_{flp}$-mNG | Km50 | This study |
| sATA525 | DH5α λpir | pSEVA231-$P_{cupE1}$-mNG | Km50 | This study |

| | | | | |
|---|---|---|---|---|
| sATA545 | DH5α λpir | pSEVA231-$P_{lecA}$-mNG | Km50 | This study |
| sATA546 | DH5α λpir | pSEVA231-$P_{lecB}$-mNG | Km50 | This study |
| sATA547 | DH5α λpir | pSEVA231-$P_{pa2566}$-mNG | Km50 | This study |
| sATA550 | DH5α λpir | pSEVA231-$P_{exsD}$-mNG | Km50 | This study |
| sATA551 | DH5α λpir | pSEVA231-$P_{lip3}$-mNG | Km50 | This study |
| sATA552 | DH5α λpir | pSEVA231-$P_{rhlA}$-mNG | Km50 | This study |
| sATA554 | DH5α λpir | pSEVA231-$P_{argA}$-mNG | Km50 | This study |
| sATA561 | DH5α λpir | pSEVA231-$P_{cupC1}$-mNG | Km50 | This study |
| sATA562 | DH5α λpir | pSEVA231-$P_{aprX}$-mNG | Km50 | This study |
| sATA563 | DH5α λpir | pSEVA231-$P_{fha1}$-mNG | Km50 | This study |
| sATA565 | DH5α λpir | pSEVA231-$P_{cupB1}$-mNG | Km50 | This study |
| sATA604 | DH5α λpir | pSEVA231-$P_{rplM}$-mNG | Km50 | This study |
| sATA605 | DH5α λpir | pSEVA231-$P_{pvdE}$-mNG | Km50 | This study |
| sATA615 | DH5α λpir | pSEVA231-$P_{pdtB}$-mNG | Km50 | This study |
| sATA620 | DH5α λpir | pSEVA231-$P_{pelA}$-mNG | Km50 | This study |
| sATA623 | DH5α λpir | pSEVA231-$P_{tssA2}$-mNG | Km50 | This study |
| sATA624 | DH5α λpir | pSEVA231-$P_{phzA1}$-mNG | Km50 | This study |
| sATA583 | DH5α λpir | pSEVA231-$P_{cupA1}$-mNG | Km50 | This study |
| sATA582 | DH5α λpir | pSEVA231-$P_{exoS}$-mNG | Km50 | This study |
| sATA703 | DH5α λpir | pSEVA231-$P_{pa2604}$-mNG | Km50 | This study |
| sATA714 | DH5α λpir | pSEVA231-$P_{rpoD}$-mNG | Km50 | This study |
| sATA707 | DH5α λpir | pSEVA231-$P_{pa1282}$-mNG | Km50 | This study |
| sATA705 | DH5α λpir | pSEVA231-$P_{xcpR}$-mNG | Km50 | This study |
| sATA700 | DH5α λpir | pSEVA231-$P_{rpsH}$-mNG | Km50 | This study |
| sATA709 | DH5α λpir | pSEVA231-$P_{pa3514}$-mNG | Km50 | This study |
| sATA783 | DH5α λpir | pSEVA231-$P_{pqsL}$-mNG | Km50 | This study |
| sATA1222 | DH5α λpir | pSEVA231-$P_{pa2384}$-mNG | Km50 | This study |
| sATA1223 | DH5α λpir | pSEVA231-$P_{pvdS}$-mNG | Km50 | This study |
| sATA1224 | DH5α λpir | pSEVA231-$P_{fptA}$-mNG | Km50 | This study |
| sATA1225 | DH5α λpir | pSEVA231-$P_{pa4570}$-mNG | Km50 | This study |
| sATA1226 | DH5α λpir | pSEVA231-$P_{prrF1}$-mNG | Km50 | This study |
| sATA1356 | DH5α λpir | pSEVA231-$P_{pyo}$-mNG | Km50 | This study |
| sATA1477 | DH5α λpir | pSEVA231-$P_{norC}$-mGL-ConvPrplM-mCh2 | Km50 | This study |
| sATA1475 | DH5α λpir | pSEVA231-$P_{hpd}$-mGL-ConvPrplM-mCh2 | Km50 | This study |
| sATA1512 | DH5α λpir | pSEVA231-$P_{flp}$-mGL-ConvPrplM-mCh2 | Km50 | This study |
| sATA2119 | DH5α λpir | pSEVA231-$P_{rhlA}$-mGL-ConvPrplM-mCh2 | Km50 | This study |
| sATA2120 | DH5α λpir | pSEVA231-$P_{exoY}$-mGL-ConvPrplM-mCh2 | Km50 | This study |
| sATA1515 | DH5α λpir | pPSV35-PprB | Gm20 | This study |
| sATA1516 | DH5α λpir | pEXG2-Δ*pprB* | Gm20 | This study |
| sATA1525 | SM10 | pEXG2-Δ*pprB* | Gm20 | This study |
| sATA253 | DH5α λpir | pEXG2-Δ*exsA* | Gm20 | This study |
| sATA263 | SM10 | pEXG2-Δ*exsA* | Gm20 | This study |
| sATA1886 | DH5α λpir | pEXG2-Δ*exsE* | Gm20 | This study |
| sATA1887 | DH5α λpir | pEXG2-Δ*exsE* | Gm20 | This study |
| sATA1870 | DH5α λpir | pSEVA231-$P_{rplM}$-mGL | Km50 | This study |
| sATA1470 | DH5α λpir | pSEVA231-$P_{rplM}$-mCh2 | Km50 | This study |

| | | | | |
|---|---|---|---|---|
| sATA2212 | DH5α λpir | pEXG2-Δ*flp* | Gm20 | This study |
| sATA2213 | SM10 | pEXG2-Δ*flp* | Gm20 | This study |
| *P. aeruginosa* PAO1 | | | | |
| sATA001 | WT (UW lineage) | - | - | [5] |
| sATA817 | WT (DSM lineage) | - | - | [6] |
| sATA647 | WT (UW lineage) | pSEVA231 | Km500 | This study |
| sATA427 | WT (UW lineage) | pPSV35 | Gm20 | [4] |
| sATA082 | WT (UW lineage) | pSEVA231-$P_{algU}$-mNG | Km500 | This study |
| sATA088 | WT (UW lineage) | pSEVA231-$P_{hpd}$-mNG | Km500 | This study |
| sATA089 | WT (UW lineage) | pSEVA231-$P_{norC}$-mNG | Km500 | This study |
| sATA114 | WT (UW lineage) | pSEVA231-$P_{fabD}$-mNG | Km500 | This study |
| sATA115 | WT (UW lineage) | pSEVA231-$P_{fhp}$-mNG | Km500 | This study |
| sATA116 | WT (UW lineage) | pSEVA231-$P_{flhA}$-mNG | Km500 | This study |
| sATA117 | WT (UW lineage) | pSEVA231-$P_{nirS}$-mNG | Km500 | This study |
| sATA118 | WT (UW lineage) | pSEVA231-$P_{nuoA}$-mNG | Km500 | This study |
| sATA119 | WT (UW lineage) | pSEVA231-$P_{pa0797}$-mNG | Km500 | This study |
| sATA120 | WT (UW lineage) | pSEVA231-$P_{recN}$-mNG | Km500 | This study |
| sATA121 | WT (UW lineage) | pSEVA231-$P_{fdnG}$-mNG | Km500 | This study |
| sATA122 | WT (UW lineage) | pSEVA231-$P_{narK1}$-mNG | Km500 | This study |
| sATA123 | WT (UW lineage) | pSEVA231-$P_{pilA}$-mNG | Km500 | This study |
| sATA124 | WT (UW lineage) | pSEVA231-$P_{aceA}$-mNG | Km500 | This study |
| sATA125 | WT (UW lineage) | pSEVA231-$P_{lldA}$-mNG | Km500 | This study |
| sATA126 | WT (UW lineage) | pSEVA231-$P_{oprO}$-mNG | Km500 | This study |
| sATA161 | WT (UW lineage) | pSEVA231-$P_{cifR}$-mNG | Km500 | This study |
| sATA162 | WT (UW lineage) | pSEVA231-$P_{pa2469}$-mNG | Km500 | This study |
| sATA163 | WT (UW lineage) | pSEVA231-$P_{aspS}$-mNG | Km500 | This study |
| sATA164 | WT (UW lineage) | pSEVA231-$P_{pchR}$-mNG | Km500 | This study |
| sATA165 | WT (UW lineage) | pSEVA231-$P_{cspA}$-mNG | Km500 | This study |
| sATA166 | WT (UW lineage) | pSEVA231-$P_{pa3754}$-mNG | Km500 | This study |
| sATA167 | WT (UW lineage) | pSEVA231-$P_{bkdA1}$-mNG | Km500 | This study |
| sATA168 | WT (UW lineage) | pSEVA231-$P_{fiuI}$-mNG | Km500 | This study |
| sATA169 | WT (UW lineage) | pSEVA231-$P_{phuR}$-mNG | Km500 | This study |
| sATA170 | WT (UW lineage) | pSEVA231-$P_{ampC}$-mNG | Km500 | This study |
| sATA171 | WT (UW lineage) | pSEVA231-$P_{fagA}$-mNG | Km500 | This study |
| sATA172 | WT (UW lineage) | pSEVA231-$P_{pa5539}$-mNG | Km500 | This study |
| sATA173 | WT (UW lineage) | pSEVA231-$P_{oprD}$-mNG | Km500 | This study |
| sATA174 | WT (UW lineage) | pSEVA231-$P_{mexA}$-mNG | Km500 | This study |
| sATA175 | WT (UW lineage) | pSEVA231-$P_{muxA}$-mNG | Km500 | This study |
| sATA176 | WT (UW lineage) | pSEVA231-$P_{lexA}$-mNG | Km500 | This study |
| sATA177 | WT (UW lineage) | pSEVA231-$P_{groES}$-mNG | Km500 | This study |
| sATA178 | WT (UW lineage) | pSEVA231-$P_{ibpA}$-mNG | Km500 | This study |
| sATA180 | WT (UW lineage) | pSEVA231-$P_{betI}$-mNG | Km500 | This study |
| sATA494 | WT (UW lineage) | pSEVA231-$P_{pchE}$-mNG | Km500 | This study |
| sATA569 | WT (UW lineage) | pSEVA231-$P_{pa0830}$-mNG | Km500 | This study |
| sATA570 | WT (UW lineage) | pSEVA231-$P_{coxB}$-mNG | Km500 | This study |
| sATA573 | WT (UW lineage) | pSEVA231-$P_{tssA1}$-mNG | Km500 | This study |
| sATA574 | WT (UW lineage) | pSEVA231-$P_{flp}$-mNG | Km500 | This study |
| sATA575 | WT (UW lineage) | pSEVA231-$P_{cupE1}$-mNG | Km500 | This study |
| sATA585 | WT (UW lineage) | pSEVA231-$P_{lecA}$-mNG | Km500 | This study |
| sATA586 | WT (UW lineage) | pSEVA231-$P_{lecB}$-mNG | Km500 | This study |

| | | | | |
|---|---|---|---|---|
| sATA587 | WT (UW lineage) | pSEVA231-$P_{pa2566}$-mNG | Km500 | This study |
| sATA588 | WT (UW lineage) | pSEVA231-$P_{exsD}$-mNG | Km500 | This study |
| sATA589 | WT (UW lineage) | pSEVA231-$P_{lip3}$-mNG | Km500 | This study |
| sATA590 | WT (UW lineage) | pSEVA231-$P_{rhlA}$-mNG | Km500 | This study |
| sATA591 | WT (UW lineage) | pSEVA231-$P_{argA}$-mNG | Km500 | This study |
| sATA592 | WT (UW lineage) | pSEVA231-$P_{cupC1}$-mNG | Km500 | This study |
| sATA593 | WT (UW lineage) | pSEVA231-$P_{aprX}$-mNG | Km500 | This study |
| sATA594 | WT (UW lineage) | pSEVA231-$P_{fha1}$-mNG | Km500 | This study |
| sATA595 | WT (UW lineage) | pSEVA231-$P_{cupB1}$-mNG | Km500 | This study |
| sATA609 | WT (UW lineage) | pSEVA231-$P_{rplM}$-mNG | Km500 | This study |
| sATA610 | WT (UW lineage) | pSEVA231-$P_{pvdE}$-mNG | Km500 | This study |
| sATA633 | WT (UW lineage) | pSEVA231-$P_{pdtB}$-mNG | Km500 | This study |
| sATA634 | WT (UW lineage) | pSEVA231-$P_{pelA}$-mNG | Km500 | This study |
| sATA635 | WT (UW lineage) | pSEVA231-$P_{tssA2}$-mNG | Km500 | This study |
| sATA636 | WT (UW lineage) | pSEVA231-$P_{phzA1}$-mNG | Km500 | This study |
| sATA663 | WT (UW lineage) | pSEVA231-$P_{cupA1}$-mNG | Km500 | This study |
| sATA664 | WT (UW lineage) | pSEVA231-$P_{exoS}$-mNG | Km500 | This study |
| sATA719 | WT (UW lineage) | pSEVA231-$P_{pa2604}$-mNG | Km500 | This study |
| sATA720 | WT (UW lineage) | pSEVA231-$P_{rpoD}$-mNG | Km500 | This study |
| sATA721 | WT (UW lineage) | pSEVA231-$P_{pa1282}$-mNG | Km500 | This study |
| sATA722 | WT (UW lineage) | pSEVA231-$P_{xcpR}$-mNG | Km500 | This study |
| sATA734 | WT (UW lineage) | pSEVA231-$P_{rpsH}$-mNG | Km500 | This study |
| sATA735 | WT (UW lineage) | pSEVA231-$P_{pa3514}$-mNG | Km500 | This study |
| sATA784 | WT (UW lineage) | pSEVA231-$P_{pqsL}$-mNG | Km500 | This study |
| sATA1227 | WT (UW lineage) | pSEVA231-$P_{pa2384}$-mNG | Km500 | This study |
| sATA1228 | WT (UW lineage) | pSEVA231-$P_{pvdS}$-mNG | Km500 | This study |
| sATA1229 | WT (UW lineage) | pSEVA231-$P_{fptA}$-mNG | Km500 | This study |
| sATA1230 | WT (UW lineage) | pSEVA231-$P_{pa4570}$-mNG | Km500 | This study |
| sATA1231 | WT (UW lineage) | pSEVA231-$P_{prrF1}$-mNG | Km500 | This study |
| sATA1381 | WT (UW lineage) | pSEVA231-$P_{pyo}$-mNG | Km500 | This study |
| sATA1483 | WT (UW lineage) | pSEVA231-$P_{norC}$-mGL-ConvP$_{rplM}$-mCh2 | Km500 | This study |
| sATA1489 | WT (UW lineage) | pSEVA231-$P_{hpd}$-mGL-ConvP$_{rplM}$-mCh2 | Km500 | This study |
| sATA1521 | WT (UW lineage) | pSEVA231-$P_{flp}$-mGL-ConvP$_{rplM}$-mCh2 | Km500 | This study |
| sATA2130 | WT (UW lineage) | pSEVA231-$P_{rhlA}$-mGL-ConvP$_{rplM}$-mCh2 | Km500 | This study |
| sATA2131 | WT (UW lineage) | pSEVA231-$P_{exoY}$-mGL-ConvP$_{rplM}$-mCh2 | Km500 | This study |
| sATA1972 | Δ*exsA* | - | - | This study |
| sATA1917 | Δ*exsE* | - | - | This study |
| sATA1539 | Δ*pprB* | - | - | This study |
| sATA1543 | Δ*pprB* | pSEVA231-$P_{flp}$-mGL-ConvP$_{rplM}$-mCh2 | Km500 | This study |
| sATA1526 | WT | pPSV35-PprB | Gm20 | This study |
| sATA1750 | WT | pPSV35-PprB + pSEVA231-$P_{flp}$-mGL-ConvP$_{rplM}$-mCh2 | Gm20 + Km500 | This study |
| sATA1946 | Δ*exsE* | pSEVA231-$P_{rplM}$-mGL | Km500 | This study |
| sATA2044 | Δ*exsA* | pSEVA231-$P_{rplM}$-mCh2 | Km500 | This study |

| | | | | |
|---|---|---|---|---|
| sATA2134 | Δ*exsA* | pSEVA231-$P_{exoY}$-mGL-ConvP$_{rplM}$-mCh2 | Km500 | This study |
| sATA2135 | Δ*exsE* | pSEVA231-$P_{exoY}$-mGL-ConvP$_{rplM}$-mCh2 | Km500 | This study |
| sATA1876 | WT | pSEVA231-$P_{rplM}$-mGL | Km500 | This study |
| sATA2215 | Δ*flp* | - | - | This study |
| sATA2243 | Δ*flp* | pSEVA231-$P_{rplM}$-mCh2 | Km500 | This study |
| sVS473 | M70639645 | - | - | [7] |
| sATA2239 | M70639645 | pSEVA231-$P_{flp}$-mGL-ConvP$_{rplM}$-mCh2 | Km500 | This study |
| sVS472 | ESP067 | - | - | [7] |
| sATA2238 | ESP067 | pSEVA231-$P_{flp}$-mGL-ConvP$_{rplM}$-mCh2 | Km500 | This study |
| sVS471 | ESP066 | - | - | [7] |
| sATA2237 | ESP066 | pSEVA231-$P_{flp}$-mGL-ConvP$_{rplM}$-mCh2 | Km500 | This study |
| sATA2230 | PSAE1903 | - | - | [7] |
| sATA2235 | PSAE1903 | pSEVA231-$P_{flp}$-mGL-ConvP$_{rplM}$-mCh2 | Km500 | This study |
| sVS476 | MHH0147 | - | - | [7] |
| sATA2240 | MHH0147 | pSEVA231-$P_{flp}$-mGL-ConvP$_{rplM}$-mCh2 | Km500 | This study |
| sATA2226 | MHH15204 | - | - | [7] |
| sATA2234 | MHH15204 | pSEVA231-$P_{flp}$-mGL-ConvP$_{rplM}$-mCh2 | Km500 | This study |
| sATA2224 | CH4757 | - | - | [7] |
| sATA2236 | CH4757 | pSEVA231-$P_{flp}$-mGL-ConvP$_{rplM}$-mCh2 | Km500 | This study |
| sATA2225 | MHH15015 | - | - | [7] |
| sATA2233 | MHH15015 | pSEVA231-$P_{flp}$-mGL-ConvP$_{rplM}$-mCh2 | Km500 | This study |
