## Supplementary Table 10 for "Subpopulations of *Pseudomonas aeruginosa* inflict distinct tissue damage during infection"

**Supplementary Table 3. Primers used in this study.**

| Primer | Sequence | Description |
|---|---|---|
| oATA002 | GAGTCCAAGACTAGTCGCCAGGG | rv primer to sequence the MCS of pSEVA plasmids |
| oATA024 | TACTCAGGAGAGCGTTCACCG | fw primer to sequence the MCS of pSEVA plasmids |
| oATA003 | ATTCGAGCTCGGTACCCGGGGATCCTGAAGCGAAGGAGGTTACGACGAAATATGGTGAGCAAAGGTGAAGAGG | fw primer to introduce mNeonGreen into BamHI restriction site of pSEVA plasmids, with an optimized RBS for 5000 expression |
| oATA004 | TCACGACGCGGCCGCAAGCTTTATTTGTACAGCTCATCC | rv primer to introduce mNeonGreen into HindIII restriction site of pSEVA plasmids, with an overlap to pSEVA |
| oATA351 | GAATTCGCGCGGCCGCGGCCTAGGcgaggcagagggcgctgactttcagg | fw primer to amplify the PrplM promoter, with an overlap to the AvrII site of pSEVA plasmids |
| oATA352 | CCTCCTCGCCCTTGCTCACCATAAATGTCTCTTAACGTAACCGTATCTAGAtcgttatagcctcaggggcc | rv primer to amplify the PrplM promoter, with an XbaI site, an RBS (strength 5000) and an overlap to the beginning of mCherry2 |
| oATA143 | ATGGTGAGCAAGGGCGAGGAGG | fw oligo to amplify mCherry2 with an ATG |
| oATA144 | TTTCGTTTTATTTGATGCCTTTAATTAATTACTTGTACAGCTCGTCCATGC | rv primer to amplify mCherry2 with STOP codon and an overlap to the PacI site of pSEVA plasmids |
| oATA852 | TATACGAGCCGGAAGCATAAATGTAAAGCAAGCTTcggcatcgttgaacgactgc | fw primer to amplify the region upstream from pprB, PA4296, to make a deletion with an overlap to clone by Gibson assembly into the HindIII restriction site |
| oATA853 | ccgcgaggccggtttgtcc | rv primer to amplify the first 7 codons of pprB, PA4296 |
| oATA854 | tggacaaaccggcctcgcggagcggagcggtggtgcactg | fw primer to amplify the last 7 codons of pprB, PA4296, with an overlap to oATA853 |
| oATA855 | GTACCGAATTCGAGCTCGAGCCCGGGGATCCTCTAGAcaacaagaattccgcggtcg | rv primer to amplify the region upstream from pprB, PA4296, to make a deletion with an overlap to clone by Gibson assembly into the XbaI restriction site |
| oATA856 | ctcgtagcccgcatcggagg | fw primer to assess the deletion of pprB, PA4296 |
| oATA857 | gcgaggtgaacagcatgtcg | rv primer to assess the deletion of pprB, PA4296 |
| oATA860 | ATGACCATGATTACGAATTCAGGAGGAAACTAGTatggacaaaccggcctcgcg | fw primer to amplify PA4296_PprB with an overlap for insertion in the EcoRI site of pPSV35 and an RBS that also contains a stop codon for the lacZa gene that starts translation in the IPTG-inducible promoter |
| oATA861 | AGGTCGACTCTAGAGGATCCtcaCTTGTCGTCATCGTCTTTGTAGTCgtgcaccaccgctccgctgc | rv primer to amplify PA4296_PprB with an overlap for insertion in the BamHI site of pPSV35 and a FLAG tag and STOP codon |
| oATA135 | GTACCGAATTCGAGCTCGAGCCCGGGGATCCTCTAGActggctggcgcgcaacgtgg | fw primer to amplify 800bp from the region upstream from exsA (PA1713), with an overlap to the XbaI site from the pEXG2 plasmid |
| oATA136 | ttatttttagcccggcattcaagagatttggctccttgc | rv primer to amplify the region upstream from exsA (PA1713) with an overlap to oATA137 |
| oATA137 | gaatgccgggctaaaaataactgacg | fw primer to amplify the region downstream from exsA (PA1713) |
| oATA138 | TATACGAGCCGGAAGCATAAATGTAAAGCAAGCTTgcagagaggttgccagaggc | rv primer to amplify 800bp from the region downstream from exsA (PA1713), with an overlap to the HindIII site from the pEXG2 plasmid |
| oATA139 | tcgctgcggtcggcatgagg | fw primer to check the deletion of exsA (PA1713) |
| oATA140 | cggcaacaaagggcccttgg | rv primer to check the deletion of exsA (PA1713) |

| | | |
|---|---|---|
| oATA1042 | GTACCGAATTCGAGCTCGAGCCCGGGGATCCTCTAGAcgcctcggtccaggtgcgc | fw primer to amplify 800bp from the region upstream from exsE, PA1711, with an overlap to the XbaI site from the pEXG2 plasmid |
| oATA1043 | cgaaatcgattcgattttcatggg | rv primer to amplify the region upstream from exsE, PA1711, including the first 7 aa |
| oATA1044 | tgaaaatcgaatcgatttcgcgcacggtccggcgcaggtg | fw primer to amplify the region downstream from exsE, PA1711, with an overlap to oATA1043 |
| oATA1045 | TATACGAGCCGGAAGCATAAATGTAAAGCAAGCTTgtttaccctgtattcgaaagttgg | rv primer to amplify 800bp from the region downstream from exsE, PA1711, with an overlap to the HindIII site from the pEXG2 plasmid |
| oATA1046 | caaggtctgggcggcggacc | fw primer to assess the deletion of exsE, PA1711 |
| oATA1047 | gtcctttcccttggtacttacg | rv primer to assess the deletion of exsE, PA1711 |
| oATA1199 | GTACCGAATTCGAGCTCGAGCCCGGGGATCCTCTAGActtcgtccacagcgatcgcc | fw primer to amplify 800bp from the region upstream from flp, PA4306, with an overlap to the XbaI site from the pEXG2 plasmid |
| oATA1200 | gaacagggtcaggttcttcattc | rv primer to amplify the region upstream from flp, PA4306, including the first 7 aa |
| oATA1201 | tgaagaacctgaccctgttcctggctccgacagcgaactgacg | fw primer to amplify the region downstream from flp, PA4306, with an overlap to oATA1200 |
| oATA1202 | TATACGAGCCGGAAGCATAAATGTAAAGCAAGCTTtcttgccggcgtcgctgacc | rv primer to amplify 800bp from the region downstream from flp, PA4306, with an overlap to the HindIII site from the pEXG2 plasmid |
| oATA1203 | cgccgggcaacaccagttgc | fw primer to assess the deletion of flp, PA4306 |
| oATA1204 | gaaacgaactggctgctacc | rv primer to assess the deletion of flp, PA4306 |
